# An H4K20me3-CTCF switch at stable TAD boundaries enforces reversible cell state transitions

**DOI:** 10.64898/2026.01.29.702647

**Authors:** Kaiser Atai, Mithun Mitra, Eliana Bohn, Richard Hu, Adriana Corvalan, David Jelinek, Laurent Voisin, Aaron Ambrus, Kirthana Sarathy, William Mason Sparks, Kenya Bonitto, Alex Dunkwu, Hilary A. Coller

## Abstract

Reversible transitions between proliferative and quiescent cell states involve widespread gene expression changes despite stable topologically associating domain (TAD) boundaries. We report a reciprocal antagonism between TAD boundary element CTCF and histone modification H4K20me3 as a central mechanism governing the proliferation-quiescence transition. Genome-wide studies and functional perturbations reveal that elevated H4K20me3 in quiescent fibroblasts displaces CTCF at specific TAD boundaries while CTCF binding predominates in proliferating cells. Increased H4K20me3 reversibly induces a compact chromatin state, elliptical nuclear morphology, and transcriptional programs associated with quiescence. Conversely, elevated CTCF binding drives open chromatin, proliferative gene expression, and cell division despite quiescence signals. Fibroblasts lacking H4K20me3 methyltransferase KMT5C/Suv4-20h2 are hyper-proliferative and KMT5C-deficient mice are larger. Our findings provide a mechanistic framework for how architectural and epigenetic regulators exchange at TAD boundaries to coordinate reversible cell state transitions, a finding with implications for organismal development and diseases of dysregulated proliferation.

**Graphical abstract:** 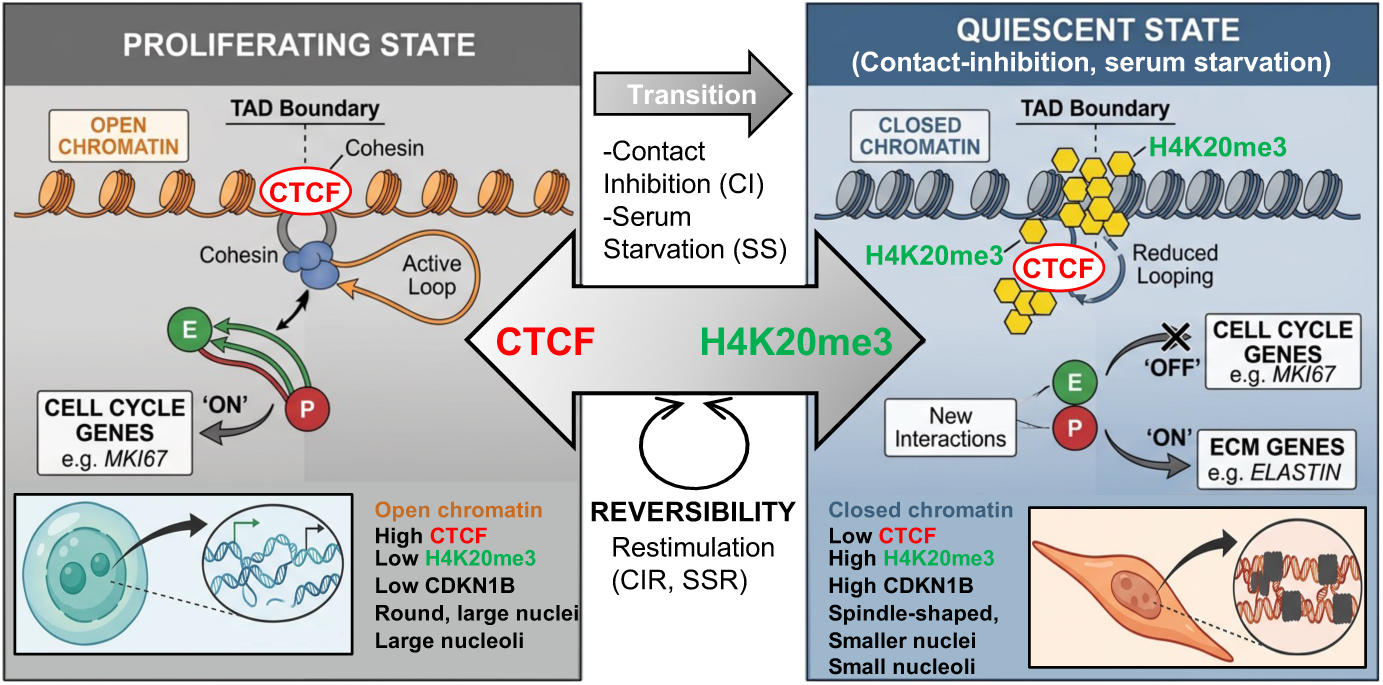

---

Chromatin is organized into topologically associating domains (TADs)^1^ that facilitate associations between regulatory elements and their target genes.^2^ Disrupting TAD boundaries can result in aberrant gene expression, with consequences for cellular function, yet, it remains unclear how gene expression and chromatin organization coordinate to regulate changes in cellular states, especially because TAD boundaries are generally invariant across cell types and conditions.^3,4^

One change in cell state that is often accompanied by altered chromatin organization is the transition between proliferation and quiescence.^5–8^ Quiescence is an actively enforced and regulated state of reversible cell-cycle arrest^9–11^ that is essential for tissue homeostasis across species. Many cell types can maintain a viable quiescent state including lymphocytes, hematopoietic stem cells, and fibroblasts.^12^ Many quiescent cells activate protective mechanisms to prevent the accumulation of waste, reduce DNA damage, and limit senescence.^9,11,13–16^ Signals that induce quiescence include serum starvation and contact inhibition.^9,17^ Many types of quiescent cells contain elevated levels of cyclin-dependent kinase inhibitor CDKN1B/p27,^18^ which plays a key role in maintaining cell cycle arrest.^18^

Fibroblasts and lymphocytes undergo changes in chromatin organization with quiescence, with most studies showing more densely packed heterochromatin in quiescent cells than their proliferating counterparts,^5–8^ while TAD boundaries are largely unchanged.^19^ Quiescent cells also possess distinct histone modification patterns,^7,20–22^ some of which are functionally important for establishing and maintaining quiescence.^7,20,22^ Quiescence-associated alterations in chromatin modification and organization are accompanied by widespread changes in gene expression in yeast^23^ and many mammalian cell types.^9,24–26^

CCCTC-binding factor (CTCF) is a highly conserved protein that binds 55,000-65,000 sites in mammalian genomes.^27^ CTCF, along with the cohesin complex, acts as an insulator by anchoring the borders between TADs.^27^ CTCF has context-dependent roles in regulating proliferation, with some studies indicating that CTCF loss reduces proliferation^28^ and other studies showing that CTCF loss results in faster growth rates.^5^

How histone modifications regulate and coordinate gene expression and chromatin organization during reversible cell state transitions remains an outstanding question in the field. Our lab previously found that the histone modification, H4K20me3, is upregulated in contact-inhibited fibroblasts.^7^ H4K20me3 is present in constitutive heterochromatin and has been associated with silencing and chromatin compaction.^29,30^ The mark represses proinflammatory genes,^31^ retrotransposons,^32^ and ribosomal RNA.^32,33^ Our previous studies showed that lowering H4K20me3 levels by knocking down KMT5C/Suv4-20h2, a H4K20me3 methyltransferase, resulted in more open chromatin.^7,30^ Here, using genomic, biochemical, and imaging-based methods, we discovered that the histone modification H4K20me3 and CTCF form a reciprocal regulatory circuit at TAD boundaries. The increase in H4K20me3 with quiescence opposes proliferation-promoting CTCF-mediated chromatin organization and gene expression in fibroblasts, thus elucidating a novel mechanism to regulate chromatin-associated changes that enables reversible transitions between cell states.

## Results

### H4K20 methyltransferase KMT5C/Suv4-20h2 and H4K20me3 are induced in quiescent fibroblasts

We performed RNA-seq on primary human dermal fibroblasts from three different donors under five conditions: (1) proliferating (P); (2a) serum starvation, (0.1% serum for 7 days); (2b) serum starved and restimulated with 10% serum; (3a) contact inhibition through full confluency for 7 days; and (3b) contact inhibited and restimulated by plating at 50% confluence (Supplementary Figure S1a). Consistent with earlier studies,^9,34^ RNA-seq revealed widespread and reproducible changes in gene expression in serum-starved and contact-inhibited cells compared with proliferating states (Fig. 1a, Supplementary Table S1). Differentially-expressed genes (DEGs, see Methods) downregulated in quiescent compared with proliferating fibroblasts (Supplementary Table S2) were enriched in Gene Ontology (GO) terms for cell cycle, DNA replication, and nucleotide metabolism (Supplementary Fig. S1b), while genes upregulated in serum-starved and contact-inhibited states were enriched for extracellular matrix organization (Supplementary Fig. S1b) Eighty-seven percent of genes downregulated and 80.4% of genes upregulated in serum-starved fibroblasts were also downregulated and upregulated in contact-inhibited fibroblasts, respectively (Supplementary Fig. S1c). The quiescence gene expression program was largely reversed when the cells were induced to re-enter the cell cycle for 24 hours (Fig. 1a).

**Figure 1.**
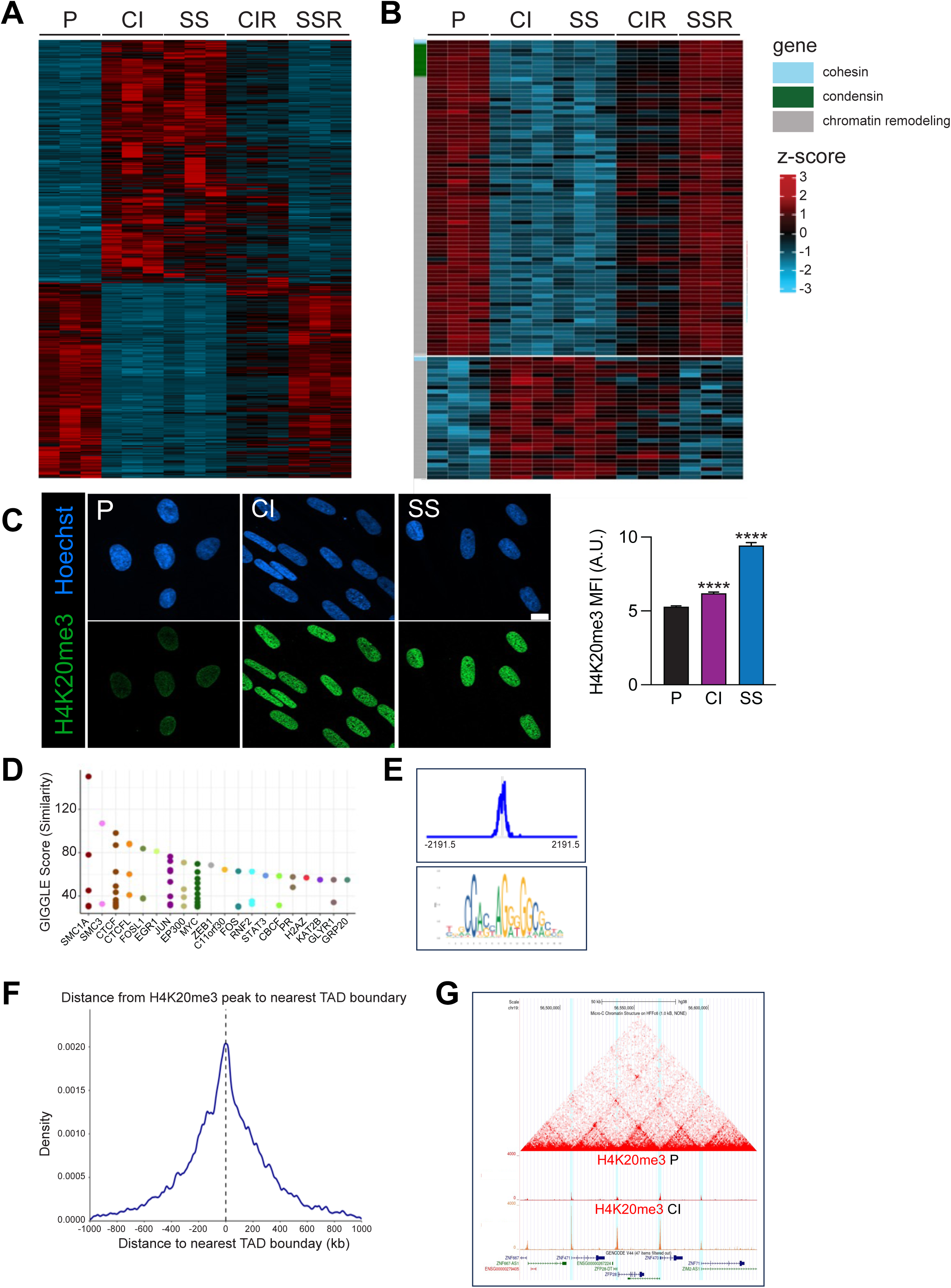
H4K20me3 and CTCF localize to the same genomic loci, which are enriched for TAD boundaries. **(A)** RNA-seq was performed on three independent lines of human fibroblasts from different donors in the indicated proliferative states. Expression values (*z*-score) for differentially expressed genes were plotted in heat map format. For all figures, P: proliferating CI: contact inhibited, SS serum starved, CIR contact inhibited and restimulated, SSR serum starved and restimulated. **(B)** Heat map for cohesin components, condensin components and differentially expressed genes in the Gene Ontology category Chromatin Remodeling (GO:0006388) with DESeq2 *p* < 0.05 and log 2-fold change >1. **(C)** Fibroblasts in the indicated proliferative conditions were analyzed with immunofluorescence for H4K20me3 levels. MFI: Mean fluorescence intensity. n = mean ± S.E.M., ANOVA. Scale bar = 10 µm. **(D)** H4K20me3 CUT&Tag peaks in CI fibroblasts were compared with publicly available datasets with Cistrome. Giggle scores for factors with the best match. **(E)** The JASPAR CTCF Motif matrix (MA0139.1) (bottom) was used to identify genomic loci predicted to be bound by CTCF. The average intensity of H4K20me3 CUT&Tag peaks across these regions (top). **(F)** Density plot of H4K20me3 CUT&Tag consensus (>2 samples) peaks across genomic regions centered at TAD boundaries. **(G)** H4K20me3 CUT&Tag data in P and CI fibroblasts with Hi-C data from fibroblasts.^38^

Among the 549 genes in the GO term for chromatin remodeling that were expressed in the fibroblasts, 38% were differentially regulated with quiescence (*p* = 1.7 x 10^-2^), with 65 upregulated and 144 downregulated (Fig. 1b and Supplementary Table S3). Consistent with higher KMT5C levels in quiescent fibroblasts (Supplementary Table S3, Fig. 1b), H4K20me3 levels were elevated in serum-starved and contact-inhibited compared with proliferating fibroblasts (Fig. 1c).

### H4K20me3 is enriched at CTCF binding sites in quiescent cells

We performed CUT&Tag to survey H4K20me3 marks in proliferating, serum-starved, and contact-inhibited fibroblasts (Supplementary Fig. S2a, b). In all samples, H4K20me3 was enriched at promoters, exons, transcription termination sites, 3’ and 5’ UTRs (Supplementary Fig. S2c), and retrotransposons (Supplementary Fig. S2d). Among genes with H4K20me3 peaks in their promoters in quiescent fibroblasts, ∼5% were upregulated with quiescence and ∼5% were downregulated. Thus, while previous studies concluded that H4K20me3 serves as a repressive mark,^31,32,35^ H4K20me3 at promoters is unlikely to be a major driver of gene repression in quiescence.

An unbiased search for the overlap between H4K20me3 CUT&Tag peaks and publicly available ChIP-seq datasets^36^ revealed that the strongest overlap was between H4K20me3 and CTCF, along with cohesins SMC1 and SMC3, which cooperate with CTCF to establish TAD boundaries ^27^ (Fig. 1d). The canonical CTCF motif was strongly enriched at sites marked by H4K20me3 (p<10^-23^). Analyzing H4K20me3 CUT&Tag data across all CTCF binding sites revealed a sharp peak of H4K20me3 intensity (Fig. 1e). Regions identified as established TAD boundaries^37^ also contained, on average, a sharp peak of H4K20me3 signal (Fig. 1f). Further, H4K20me3 peaks were visible at TAD boundaries defined by Hi-C fibroblast data^38^ (Fig. 1g).

### CTCF levels decline with quiescence

CTCF transcript levels were lower in contact-inhibited relative to proliferating fibroblasts and increased when either serum-starved or contact-inhibited fibroblasts were stimulated to re-enter the cell cycle. Consistent with this finding, we observed a 68.5% reduction in nuclear CTCF protein (Fig. 2a), along with a ∼61.2% reduction in whole-cell CTCF protein and a ∼48.6% reduction in chromatin bound-CTCF (Fig. 2b). CTCF protein abundance increased to levels like those in proliferating fibroblasts when either serum-starved or contact-inhibited fibroblasts were restimulated (Fig. 2a), demonstrating that low CTCF levels are a reversible characteristic of quiescent fibroblasts.

**Figure 2.**
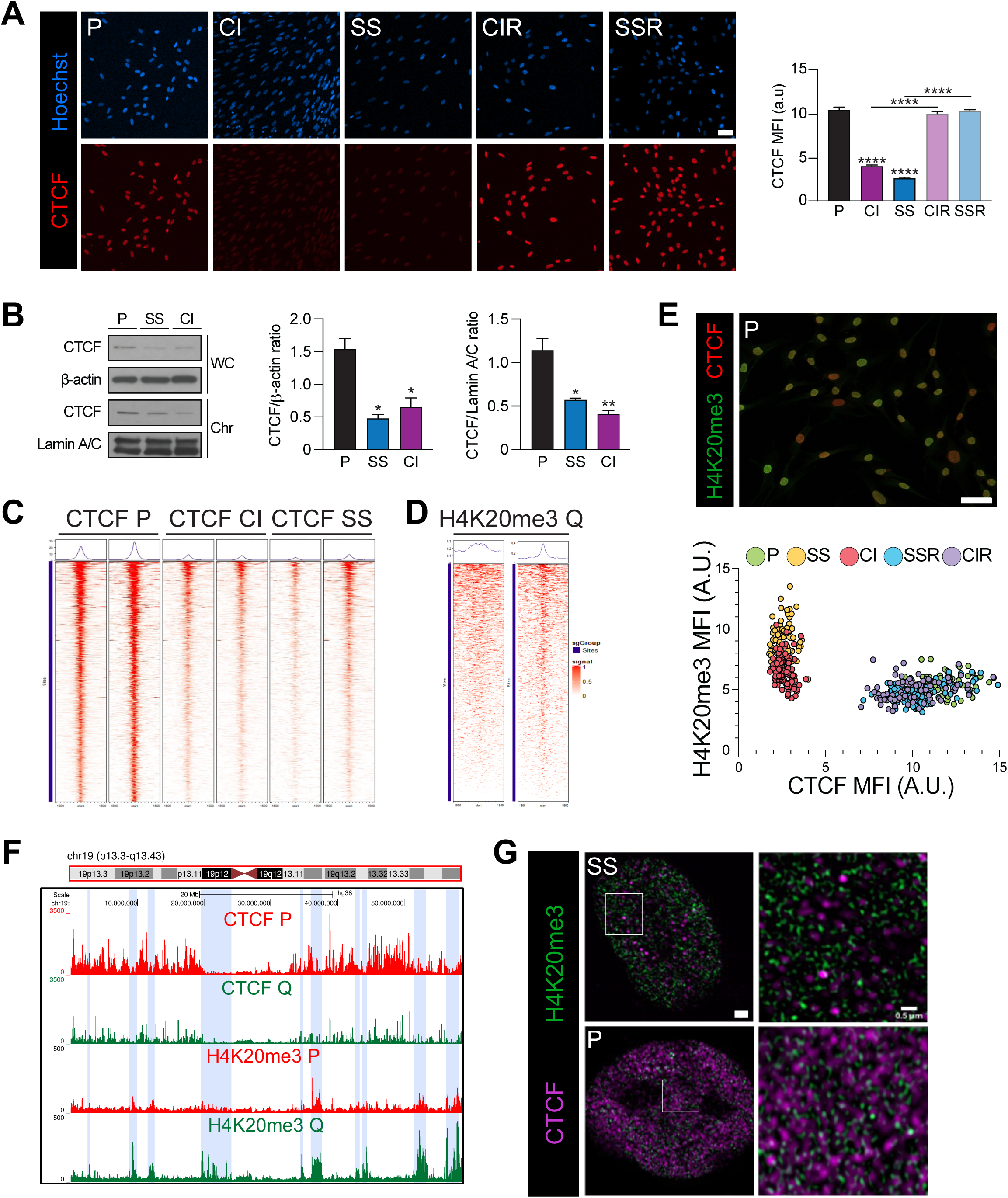
Quiescent fibroblasts contain H4K20me3-enriched chromatin regions while proliferating cells contain enhanced CTCF-bound chromatin. **(A)** Fibroblasts in different proliferative states were analyzed for CTCF with immunofluorescence and stained with Hoescht for nuclei. Arbitrary units: a.u. n = 100-260. Mean ± SEM, ANOVA. Scale bar = 10 µm. **(B)** Immunoblots were performed on whole-cell CTCF with β-actin as a loading control and for chromatin-bound CTCF with lamin A/C as a loading control. n = 3. Mean ± SEM, ANOVA. **(C)** Trackplots of CTCF CUT&Tag data for two replicates each of fibroblasts in the indicated proliferative states for peaks differentially present in proliferating and quiescent (SS and CI) cells. **(D)** Trackplots of H4K20me3 CUT&Tag data across sites with CTCF peaks that do not change between proliferating and quiescent fibroblasts (left) and sites with CTCF peaks that are reduced in quiescent fibroblasts (right). **(E)** Fibroblasts in the indicated proliferative states were analyzed with dual color immunofluorescence for CTCF and H4K20me3. n = 530 total cells. **(F)** Genome Browser views of CUT&Tag data for chromosome 19 for CTCF and H4K20me3 in proliferating and quiescent (CI and SS) fibroblasts. **(G)** Dual-color immunofluorescence for CTCF (purple) and H4K20me3 (green) was performed on P and SS fibroblasts with super-resolution microscopy. Scale bar = 1 µm (left) and 0.5 µm (right). **(H)** Chromosome-level illustration of the density of differentially expressed genes with quiescence (top chromosome, color scale) and the genomic regions with differential CTCF CUT&Tag peaks (bottom chromosome, red).

### Quiescence reduces CTCF binding at >1000 genomic loci with a concomitant increase in H4K20me3

To compare the genomic distribution of H4K20me3 and CTCF, we performed CUT&Tag for CTCF in six samples of proliferating, serum-starved, and contact-inhibited fibroblasts (Supplementary Fig. S2a, S2b). Across all samples, CTCF was enriched at promoters, exons, transcription start sites, 5’ UTRs (Supplementary Fig. S3a), and retrotransposons (Supplementary Fig. S3b). There were 1,149 sites with significant loss of CTCF binding in quiescent compared to proliferating cells (Fig. 2c, Supplementary Fig. S3c), while only 14 sites were more strongly bound by CTCF in quiescent than proliferating fibroblasts (Supplementary Fig. S3c). Sites at which CTCF was reduced with quiescence on average contained a robust H4K20me3 signal in quiescent cells (Fig. 2d), while no clear H4K20me3 signal was observed at CTCF peaks that were unchanged with quiescence (Fig. 2d). Sites with reduced CTCF binding were more likely to be located within 100 kb of a TAD boundary than all CTCF sites (38.9% vs 42.9%, +4.6 percentage points, 95% confidence interval: 0.9%, 8.8%). 11,266 loci with both CTCF and H4K20me3 peaks represented 38.3% of all CTCF peaks and 31.0% of all H4K20me3 peaks. Our results suggest a reciprocal relationship in which H4K20me3 and CTCF can associate with the same genomic loci, but each predominates in a different proliferative condition.

### H4K20me3 increases uniformly and reversibly with quiescence

Quantifying CTCF and H4K20me3 with dual-color immunofluorescence in individual cells within each of the five proliferative conditions revealed relatively homogeneous staining among the cells in each population. Proliferating and quiescent cells were clearly distinguishable from each other with proliferating and restimulated cells containing similarly high levels of CTCF and low levels of H4K20me3 (Fig. 2e). Therefore, the reciprocal pattern observed in proliferating versus quiescent cells is not confounded by heterogeneity in CTCF and H4K20me3 levels among cells within a population in a single proliferative state.

### CTCF and H4K20me3 occupy distinct chromatin regions and subnuclear territories

Viewing CTCF and H4K20me3 CUT&Tag data in proliferating and quiescent fibroblasts across an entire chromosome revealed large-scale genomic regions enriched for H4K20me3 with low CTCF (Fig. 2f, blue shading), while distinct regions contained more bound CTCF with reduced H4K20me3 levels (Fig. 2f, no shading). As cells transitioned between proliferation and quiescence, the broad genomic regions that were rich in either CTCF or H4K20me3 did not change, suggesting maintenance of a consistent pattern of large-scale chromatin organization between proliferating and quiescent cells. During proliferation, the CTCF-rich regions contained even more bound CTCF, while in quiescent conditions, the H4K20me3-rich regions accumulated additional H4K20me3. These data suggest a model for transitions between states in which large-scale boundaries remain intact, but the decoration with factors or marks within boundaries shifts between higher and lower intensity states.

With immunofluorescence and super-resolution microscopy, we found CTCF-rich punctate nuclear territories and distinct, mostly non-overlapping, regions that were H4K20me3-rich (Fig. 2g). Proliferating nuclei contained more CTCF-rich regions and fewer H4K20me3-rich regions, while serum-starved nuclei contained more H4K20me3-rich regions and fewer CTCF-rich regions (Fig. 2g). The H4K20me3-rich puncta had diameters of ∼0.05 µm and were more condensed and compact than the larger, more diffuse CTCF-rich regions, with diameters of ∼0.2 µm. The results are consistent with separate chromosomal regions and associated spatial territories within the nucleus containing largely either CTCF-associated or H4K20me3-marked chromatin. A transition to quiescence was associated with a shift to more H4K20me3-rich condensed foci and fewer CTCF-enriched diffuse foci, consistent with tighter compaction in quiescent fibroblasts.^7^

### TADs flanked by sites of lower CTCF binding in quiescence are enriched for genes regulated with quiescence

We asked whether the shift between CTCF-associated and H4K20me3-modified chromatin is associated with gene expression changes that occur with quiescence (Fig. 1a). We observed a high degree of overlap between genomic loci with reduced CTCF binding in quiescence and quiescence DEGs (Fig. 2h). We compared the expression of genes within TADs whose boundaries were within 50 kb of a CTCF peak downregulated in quiescent conditions (295 TADs) to gene expression in TADs with no change in CTCF (2,676 TADs). TADs with boundaries close to CTCF peaks with reduced occupancy showed a significant enrichment for DEGs (Wilcoxon *p* = 0.00878). The enrichment was also significant when enrichment was measured as the number of differentially expressed genes divided by the total number of genes in a TAD (permutation test with 10⁶ iterations, *p* = 0.0399). Among 2,937 DEGs, 641 genes (21.8%) were present in TADs demarcated by CTCF downregulated sites. These data support a model in which loss of CTCF binding at TAD boundaries contributes to transcriptional reprogramming with quiescence.

### H4K20me3 depletion increases chromatin-bound CTCF

To test for an antagonistic relationship between H4K20me3 and CTCF, we depleted H4K20 multimethylated marks with the pharmacological SUV4-20H inhibitor A-196, a potent and selective noncompetitive inhibitor of H4K20 methyltransferases Suv4-20h1/KMT5B and Suv4-20h2/KMT5C that does not interfere with either enzyme’s noncatalytic functions.^39,40^ We confirmed that A-196 treatment strongly decreased H4K20me3 levels in proliferating and serum-starved fibroblasts (Fig 3a, Supplementary Figure S4a). Nuclear CTCF increased significantly in both proliferating and serum-starved fibroblasts when multimethylated H4K20 was depleted with A-196 treatment (Fig. 3a). Treatment with A-196 resulted in a 2-fold increase in the levels of chromatin-bound CTCF in proliferating and serum-starved fibroblasts without affecting total CTCF levels (Fig. 3b), showing that multimethylated H4K20 can antagonize CTCF binding and supporting an inverse relationship between CTCF and H4K20me3 for chromatin association.

**Figure 3.**
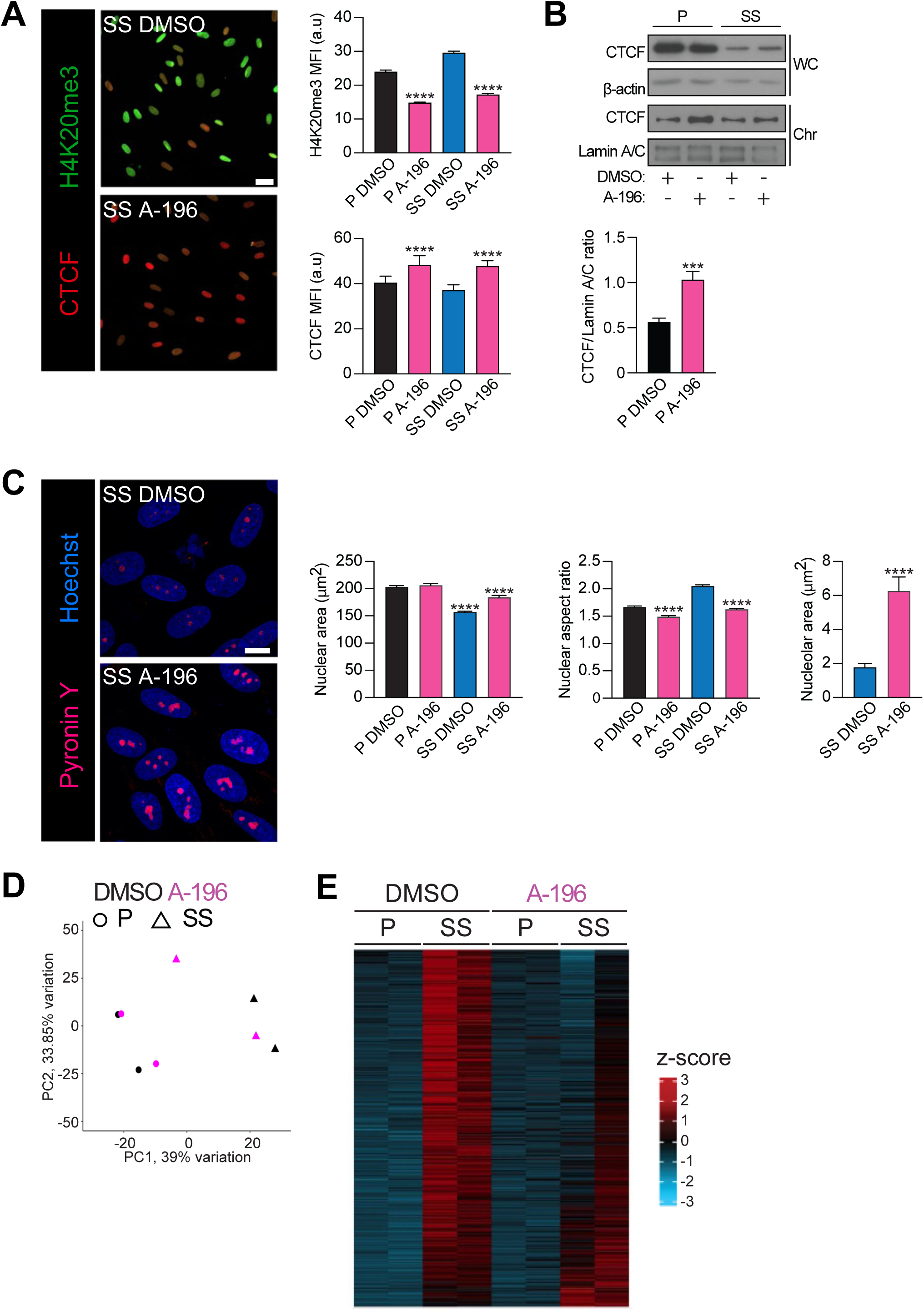
Preventing H4K20 multimethylation increases chromatin-bound CTCF, induces larger and rounder nuclei, and inhibits quiescence gene expression. **(A)** P and SS fibroblasts were treated with A-196 or DMSO as vehicle control and immunostained for H4K20me3 (green) and CTCF (red). n = 250-370. Mean ± SEM, ANOVA. Scale bar = 100 µm. **(B)** P and SS fibroblasts were treated with A196 or DMSO as vehicle control. Immunoblots were performed on whole-cell CTCF with β-actin as a loading control and for chromatin-bound CTCF with lamin A/C as a loading control. n = 2-5. Mean ± SEM, two-tailed, unpaired t-test. **(C)** P and SS fibroblasts were treated with A-196 or DMSO as vehicle control. Samples were stained with pyronin Y and Hoescht. Nuclear area and aspect ratio were determined with images of Hoescht staining. n = 255-373. Mean ± SEM, t-test. Nucleolar area (µm^2^) was determined as the pyronin Y-positive region. n = 15-25. Scale bar = 10 µm. **(D)** Principal component analysis of RNA-seq data for fibroblasts from two different cell lines in P or SS conditions treated with A-196 or DMSO as vehicle control. **(E)** Heat map of RNA-seq on 2 independent lines of human fibroblasts from different donors in P and SS conditions after treatment with A-196 or DMSO as vehicle control. Z-scores for differentially expressed genes are shown in heat map format.

### H4K20me3 depletion recapitulates morphological changes associated with proliferation

We asked whether the increased SUV4-20H activity with quiescence is important for the morphological changes that accompany cell quiescence. When fibroblasts are induced into quiescence by either serum starvation or contact inhibition, they protruded longer extensions and adapted a more extended shape (Supplementary Fig. S1a). In serum-starved and contact-inhibited fibroblasts, nuclear area decreased and nuclear aspect ratio increased compared with proliferating fibroblasts (Supplementary Fig. S5a), while proliferating fibroblasts had a larger nuclear area and lower nuclear aspect ratio (Supplementary Fig. S5a). Stimulating the fibroblasts to re-enter the cell cycle reversed these quiescence features (Supplementary Fig. S5a). 3D nuclear reconstruction confirmed smaller nuclear volumes in quiescent than proliferating fibroblasts (Supplementary Fig. S5b). Flow sorting for only G1 cells in the proliferating population confirmed that the smaller size of nuclei in quiescent fibroblasts persisted when proliferating cells in S and G2/M were excluded (Supplementary Fig. S5c). Quiescence was also associated with reduced nucleolar area (Supplementary Fig. S5d). Serum-starved fibroblasts treated with A-196 had significantly larger nuclei than vehicle DMSO-treated controls (Fig. 3c). Further, the increase in nuclear aspect ratio (long to short axis) when fibroblasts entered quiescence by serum starvation was reversed by A-196 treatment (Fig. 3c). Treatment with A-196 also strongly increased nucleolar area in serum-starved fibroblasts (Fig. 3c). Thus, impeding H4K20me3 deposition as cells enter quiescence prevented the adoption of a smaller, more elongated nucleus, demonstrating the importance of H4K20 methylation for nuclear rearrangement.

### H4K20me3 depletion regulates expression of quiescence genes

To explore the impact of H4K20 methylation on gene expression, we performed RNA-seq on proliferating and serum-starved fibroblasts treated with A-196 or DMSO. Genes downregulated by A-196 treatment were enriched for GO terms for extracellular matrix organization and extracellular structure organization (Supplementary Fig. S4b), classes of genes that are also regulated as fibroblasts enter quiescence (Supplementary Fig. S1b).^15,41^ Fifty-two out of 266 genes upregulated by A-196 treatment (19.5%) overlapped with DEGs downregulated with serum starvation, significantly more than expected by chance (hypergeometric test *p*=3.34 x 10^-43^). Further, A-196 treatment shifted serum-starved fibroblasts toward a proliferative gene expression profile (Fig. 3d) consistent with genes induced by serum starvation being less strongly upregulated when fibroblasts were serum-starved while treated with A-196 (Fig. 3e).

### Reducing CTCF levels increases H4K20me3

To test whether the relationship between H4K20me3 and CTCF is reciprocal, we investigated the impact of RNAi-mediated reduction in CTCF levels on H4K20me3. Transfecting fibroblasts with a combination of two different siRNAs targeting CTCF resulted in an 88% decrease in whole-cell CTCF levels (Fig. 4a and Supplementary Fig. S6a). Proliferating fibroblasts transfected with siRNAs targeting CTCF contained ∼50% higher levels of H4K20me3 than fibroblasts transfected with a control siRNA (Fig. 4b and Supplementary Fig. S6b).

**Figure 4.**
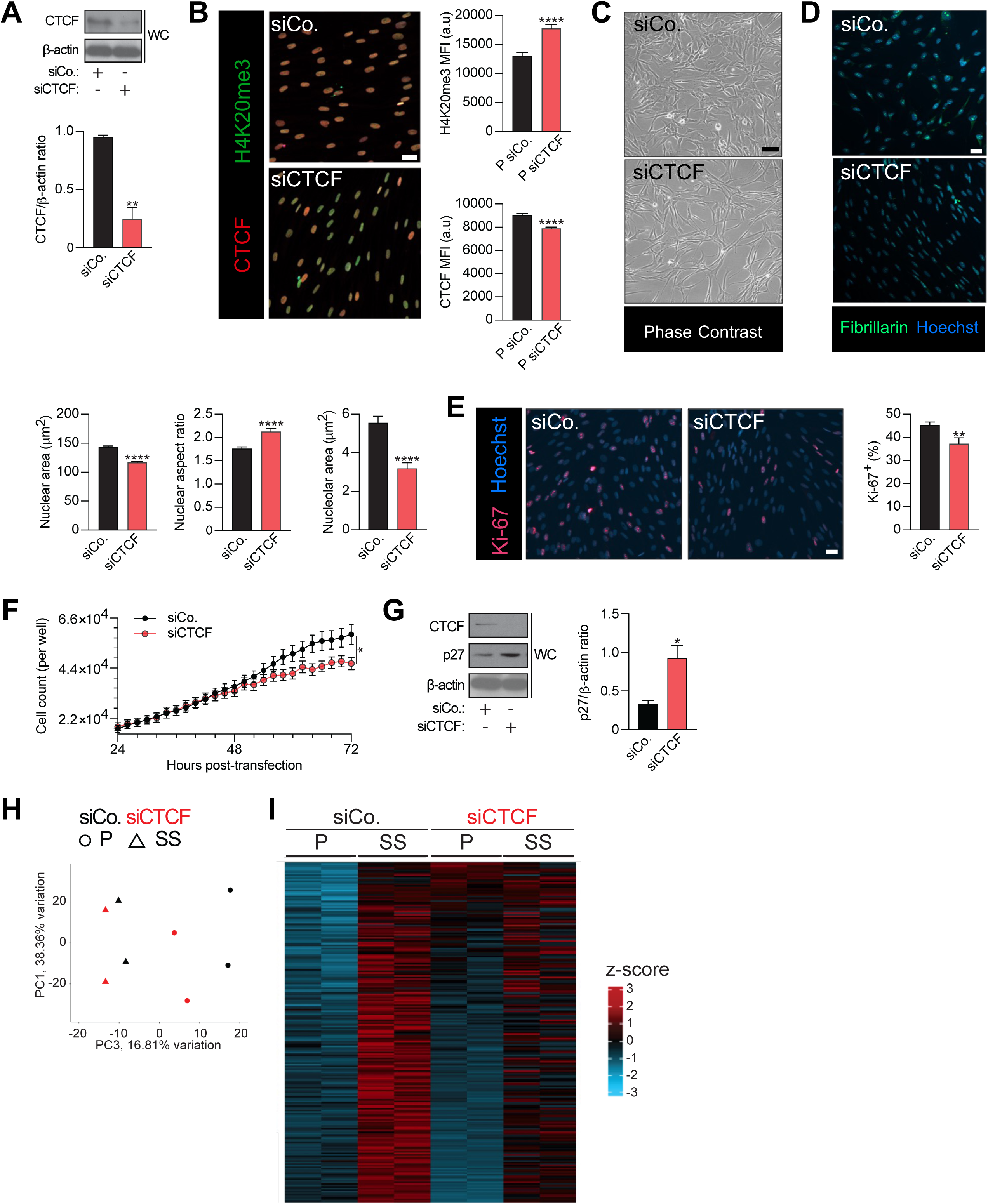
CTCF knockdown results in elevated H4K20me3, quiescence-like cell morphology, quiescence markers, reduced proliferation, and a shift toward quiescence gene expression patterns. **(A)** Fibroblasts were transfected with a control siRNA or an siRNA that targets CTCF. Protein lysates from two days after transfection were analyzed with immunoblotting for CTCF levels with β-actin as a loading control. n = 3, mean ±SEM. Complete western in Supplementary Figure S6A. **(B)** Proliferating fibroblasts transfected with a control siRNA or siRNAs against CTCF were immunostained for CTCF (red) or H4K20me3 (green). n = 170-209. Mean ± SEM, t-test. Scale bar = 100 µm. **(C)** Proliferating fibroblasts transfected with a control siRNA or a combination of two siRNAs targeting CTCF. **(D)** Proliferating fibroblasts transfected with a control siRNA or a combination of two siRNAs against CTCF were stained with Hoescht for DNA and immunofluorescence for fibrillarin. Nuclear area, nuclear aspect ratio (ratio of long to short axis), and nucleolar area were plotted. n = 490-770 cells. Mean ± SEM, t-test. **(E)** Proliferating fibroblasts were labeled with immunofluorescence for Ki-67 with Hoescht. n = 470-600 cells. Mean ± SEM, t-test. **(F)** Fibroblasts were transfected with a combination of two siRNAs against CTCF or a control siRNA and the number of cells per well was monitored with an Incucyte instrument. CTCF levels were reduced approximately 48 hours after transfection. n = 54 images every 2 hours across 6 wells. Mean ± SEM, multiple t-tests. **(G)** Protein lysates from fibroblasts transfected with a control siRNA or siRNAs that target CTCF were analyzed with immunoblotting for CTCF and p27 with β-actin as a loading control. n = 4. Mean ± SEM, t-test. **(H)** PCA plot of RNA-seq data of P and SS fibroblasts from two different donors transfected with a control siRNA or siRNAs that target CTCF. **(I)** RNA-seq was performed on 2 independent lines of human fibroblasts in P and SS conditions after transfection with siRNAs against CTCF or a control siRNA. Z-scores for differentially expressed genes were plotted in heat map format.

Given that depletion of H4K20me3 resulted in a shift in nuclear size and shape towards proliferation, we asked whether depleting CTCF would phenocopy attributes of quiescent fibroblasts. Proliferating fibroblasts with reduced CTCF adopted a distinctive morphology with thinner cell bodies and longer protrusions than control fibroblasts (Fig. 4c), similar to the appearance of serum-starved fibroblasts (Supplementary Fig S1a). CTCF depletion also resulted in a significant reduction in nuclear area (Fig. 4d), decreased nucleolar area (Fig. 4d), and more elongated nuclei (Fig. 4d, Supplementary Fig. S6c). Therefore, reducing CTCF levels results in cell morphology changes that phenocopy quiescence morphology, suggesting that reduced CTCF may mediate some quiescence characteristics.

### CTCF depletion inhibits proliferation and promotes quiescence

Consistent with CTCF depletion inducing quiescence features, CTCF knockdown resulted in a reduced fraction of Ki-67 positive cells in a proliferative culture (Fig 4e), a slowing of the proliferation rate (Fig. 4f), and upregulation of the quiescence marker CDK inhibitor CDKN1B/p27 (Fig. 4g). RNA-seq on proliferating fibroblasts transfected with siRNAs against CTCF or a control siRNA revealed that genes downregulated by both serum starvation and CTCF depletion were enriched in GO categories for cytokinesis, mitotic cell cycle, and nucleotide biosynthetic process (Supplementary Fig. 7a), while genes upregulated by both quiescence and CTCF knockdown were enriched for GO categories for extracellular matrix organization (Supplementary Fig. 7b). Genes differentially expressed by serum starvation significantly overlapped with genes regulated by CTCF knockdown (1,701 siCTCF DEGs, 3,502 serum-starved DEGs, 575 overlapping genes, hypergeometric test *p* = 2.33x10^-43^), consistent with CTCF loss partially recapitulating gene expression changes associated with quiescence. CTCF-depleted proliferating cells more closely resembled serum-starved cells transcriptionally (Fig. 4i). Thus, depleting CTCF shifts cells toward a quiescent phenotype based on multiple functional attributes, including nuclear shape, cell cycle rate, gene expression profile, proliferation markers, and p27 upregulation.

### CTCF overexpression results in larger, rounder nuclei, a shift toward proliferation-associated gene expression, and resistance to cell cycle exit

To test whether CTCF overexpression is sufficient to modulate H4K20me3 levels and proliferative state, we transduced fibroblasts with lentiviral vectors that overexpress CTCF or mCherry as a control (Supplementary Fig. S8a). When CTCF-overexpressing fibroblasts were serum-starved, H4K20me3 levels were significantly lower than in controls (Fig. 5a, Supplementary Fig. S8b), further supporting antagonistic crosstalk between CTCF and H4K20me3. Serum-starved fibroblasts overexpressing CTCF had larger and rounder nuclei and larger nucleoli than control fibroblasts (Fig. 5a, b), mimicking the changes observed as fibroblasts transition from quiescence to proliferation (Supplementary Fig. S5). CDKN1B/p27 was less abundant in serum-starved fibroblasts overexpressing CTCF than serum-starved control fibroblasts (Fig. 5c), demonstrating that quiescence induction was impeded by excess CTCF. The fraction of Ki-67-positive cells among CTCF-overexpressing fibroblasts was higher than controls on days 1, 4, and 7 after serum was withdrawn (Fig. 5d). Even after 7 days of serum starvation, 11% of CTCF-overexpressing fibroblasts remained Ki-67-positive, compared with just 2% of control fibroblasts (Fig. 5d).

**Figure 5.**
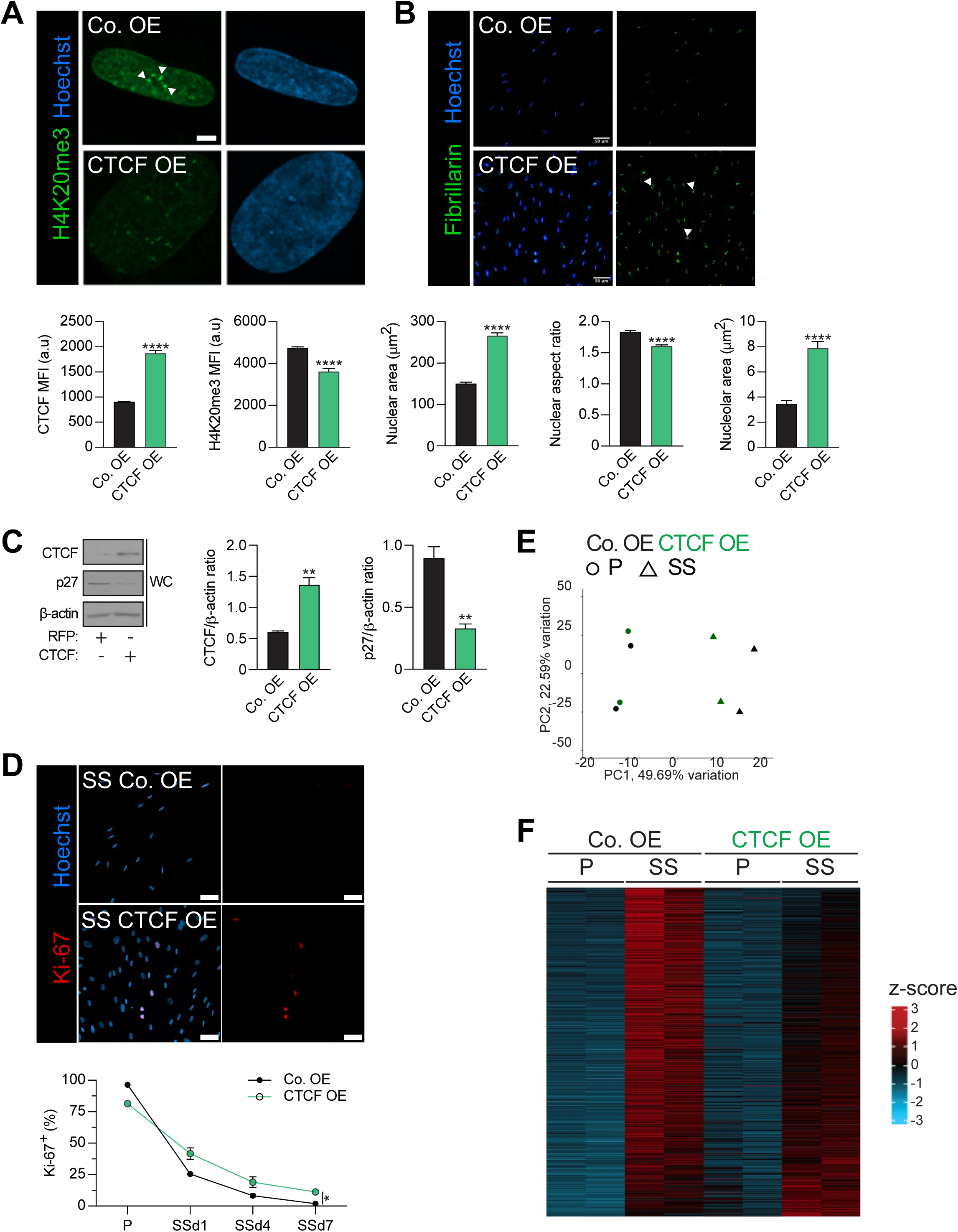
CTCF overexpression reduces H4K20me3 levels and allows fibroblasts to resist serum starvation-induced cell cycle arrest, quiescence markers, and gene expression changes. **(A)** Fibroblasts overexpressing CTCF or a control protein were immunostained for CTCF and H4K20me3 and stained with Hoescht. n = 315-670 cells for MFI, n = 350-960 for nuclear area and nuclear aspect ratio. Mean ± SEM, t-test. Scale bar = 5 µm. **(B)** Serum-starved fibroblasts overexpressing CTCF or a control protein were immunostained for fibrillarin and stained with Hoescht. n = 43-98. Mean ± SEM, t-test. Scale bar = 50 µm. **(C)** Protein lysates from fibroblasts overexpressing CTCF or a control protein were analyzed with immunoblotting for CTCF and CKI CDKN1B/p27. **(D)** Fibroblasts overexpressing CTCF or a control protein were serum-starved and collected on days 1, 4 and 7 after serum withdrawal. Fibroblasts were immunostained for Ki-67 and stained with Hoescht. n = 925 cells. Mean ± SEM, multiple t-tests. All data points are significant. Scale bar = 50 µm. **(E)** Principal component analysis of RNA-seq data for fibroblasts from two different cell lines in P or SS conditions overexpressing CTCF or control protein. **(F)** RNA-seq was performed on P and SS fibroblasts transduced with lentiviral vectors overexpressing CTCF or a control protein. Z-scores for quiescence differentially expressed genes are shown in heat map format.

RNA-seq data showed that CTCF overexpression shifted the transcriptome of serum-starved fibroblasts towards a proliferative pattern (Fig. 5e), while gene expression patterns in serum-starved fibroblasts show that CTCF overexpression dampens the induction of quiescence DEGs (Fig. 5f), consistent with CTCF downregulation playing a critical role in the establishment of a quiescence gene expression program. Thus, CTCF acts as a dominant determinant of proliferative identity within this regulatory circuit.

### *Mki67* is flanked by CTCF and H4K20me3-bound sites

Our CTCF CUT&Tag data show that the gene encoding the Ki-67 protein, *Mki67*, is flanked by CTCF-bound sites (Fig. 6a, yellow regions) and does not have a strong CTCF binding site at its promoter. Fibroblast Hi-C data^38^ suggests the CTCF-binding chromosomal regions anchor a chromatin loop (Fig. 6a, yellow regions). CTCF levels were higher in proliferating than quiescent cells at both *Mki67*-flanking sites (Fig. 6a). H4K20me3 peaks were at these same loci were particularly high in quiescent cells (Fig. 6a). *Mki67* expression levels were 7.2-fold lower in serum-starved compared with proliferating fibroblasts (Fig. 6c). With CTCF overexpression, *Mki67* expression levels increased 1.4-fold, while siRNA-mediated depletion of CTCF resulted in a 1.6-fold decrease in expression relative to control fibroblasts (Fig. 6c). The results suggest *Mki67* expression is induced in proliferating fibroblasts in part through CTCF-mediated chromatin rearrangements.

**Figure 6.**
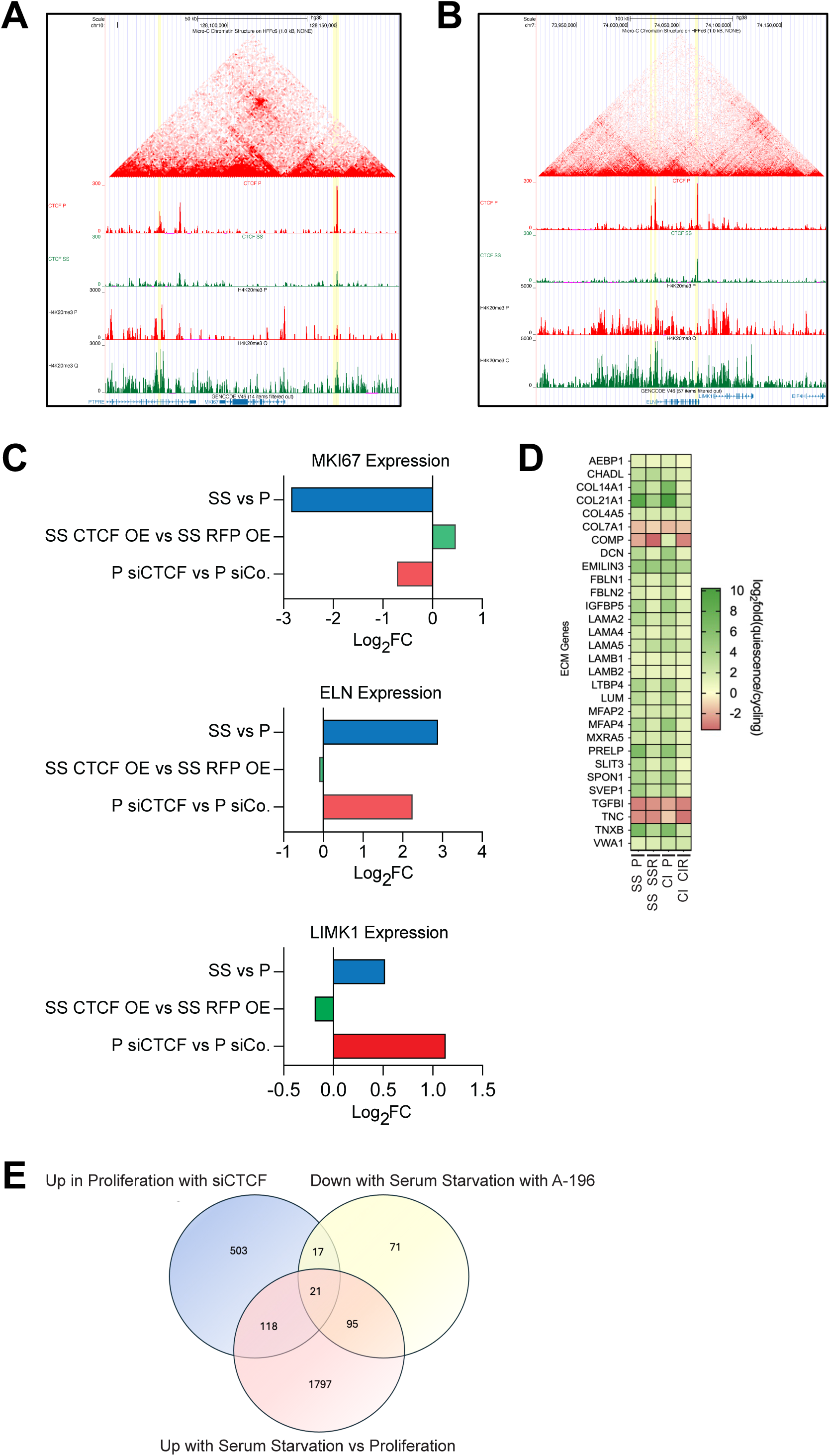
***Mki67* and *Elastin* are flanked by differential CTCF binding sites and regulated by CTCF expression (A, B)** UCSC Genome Browser views of HiC data, along with CTCF and H4K20me3 CUT&Tag data in P and CI fibroblasts near *Mki67* (A) or *Elastin* and *Limk* (B). Yellow highlights genomic loci with differential CTCF binding and H4K20me3 marks. **(C)** Log_2_ fold change RNA-seq expression data are plotted for *Mki67*, *Elastin*, and *Limk1* for P vs SS fibroblasts, CTCF vs control overexpression in SS fibroblasts, and siCTCF vs siControl in P fibroblasts. **(D)** Heat map of relative gene expression levels for genes in the GO category for extracellular matrix that are also differentially expressed genes with quiescence. Log_2_ fold change is shown for quiescence versus proliferating/restimulated. **(E)** RNA-seq was performed on SS and P fibroblasts, A196- or DMSO-treated fibroblasts, and fibroblasts transfected with siRNAs to CTCF or a control siRNA. A Venn diagram shows the extent of overlap among genes upregulated with serum starvation, induced with siCTCF, and downregulated with A-196 treatment out of 16,915 genes expressed in all conditions. All overlaps have hypergeometric *p* values of 10^-17^ or lower. The three-way intersection is 22-fold enriched versus independence expectation.

### Genes associated with the extracellular matrix are regulated in response to quiescence, CTCF overexpression, and H4K20me3 depletion

While CTCF binding at a loop anchor can induce gene expression by facilitating promoter-enhancer interactions, loss of CTCF binding, as occurs during quiescence, can affect gene expression by relieving repression from silencing elements within a TAD or by weakening loop formation.^42^ In our data, 26 out of 30 ExtraCellular Matrix (ECM) components that were regulated with quiescence were upregulated (Fig. 6d). Genes upregulated with serum starvation, downregulated with A-196 (Supplementary Figure S4b), and upregulated with CTCF knockdown (Supplementary Figure S7b) were all enriched for the GO term “extracellular matrix” and highly significantly overlapped (22-fold enrichment, hypergeometric *p* = 4.2 x 10^-17^) (Fig. 6e).

As an example of an ECM gene, the gene encoding Elastin (*Eln*) is flanked by genomic loci (Fig. 6b yellow regions) that are anchor sites for a chromatin loop based on Hi-C in proliferating fibroblasts (Fig. 6b).^38^ Both sites have greater CTCF binding in proliferating fibroblasts and enhanced H4K20me3 in quiescent cells (Fig. 6b). *Eln* expression levels were 7.4-fold higher in serum-starved fibroblasts and 4.8-fold-higher with CTCF knockdown (Fig. 6c). These findings suggest that CTCF regulates gene expression through modulating the regulatory effects of existing TAD architecture in ways that can increase or decrease gene expression.

### Chromatin is more open and CTCF is upregulated in wound-healing fibroblasts

To test whether fibroblasts in animals follow patterns like cultured fibroblasts, we compared proliferative fibroblasts adjacent to dermal mouse wounds with quiescent fibroblasts in unwounded mouse skin.^24^ We introduced incisional wounds into the flanks of C57BL/6 mice and used transmission electron microscopy to image the dermal fibroblasts in unwounded skin and wound samples collected 1 day after the wounds were introduced. Similar to quiescent cultured human fibroblasts that contain more condensed chromatin than proliferating fibroblasts,^7^ the fraction of the nuclear area composed of electron-dense heterochromatin was elevated in quiescent fibroblasts from unwounded skin compared with proliferating fibroblasts in wound-adjacent skin (Fig. 7a).

**Figure 7.**
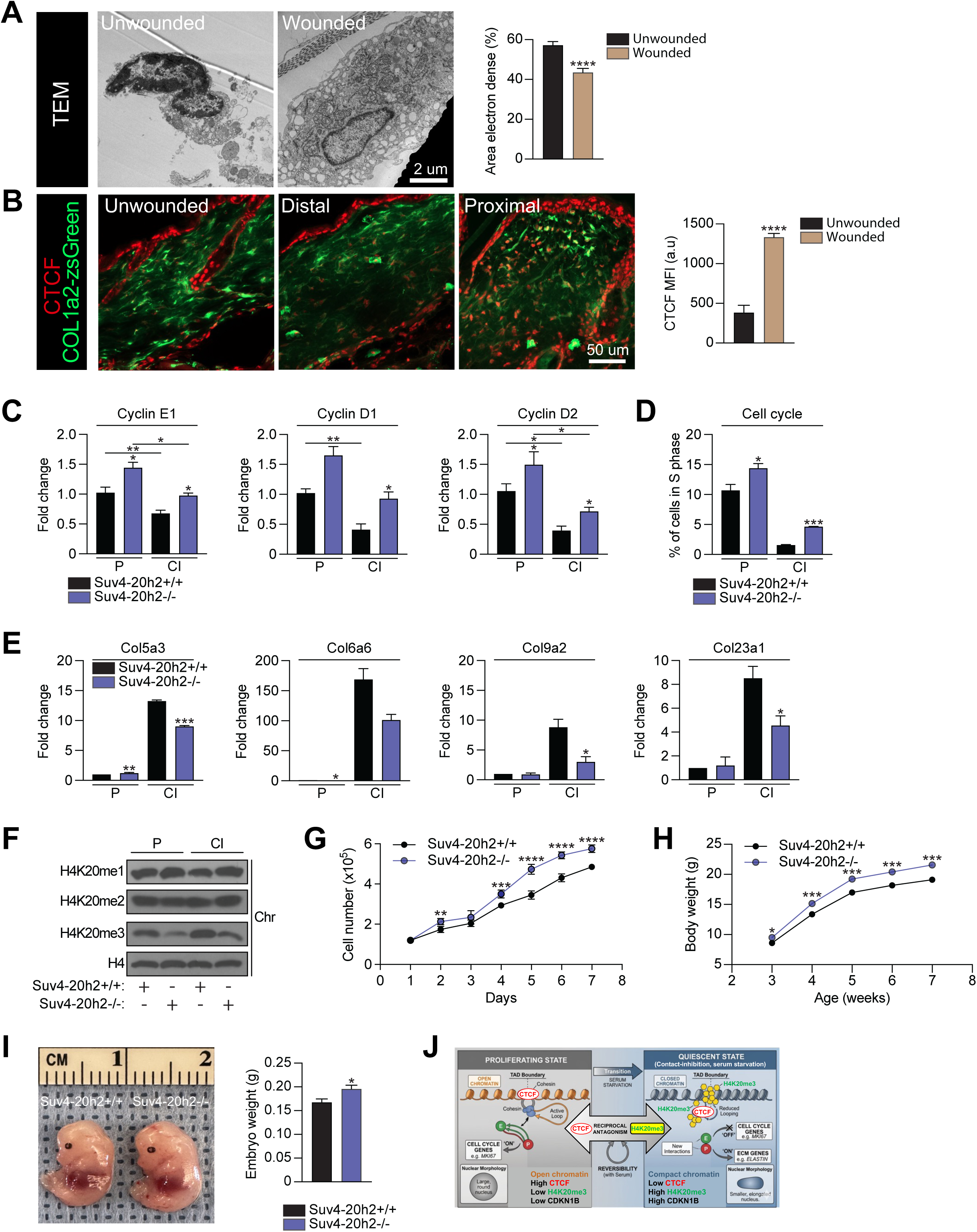
CTCF is upregulated in the fibroblast wound healing response, *Suv4-20h2*-deficient MEFs are more proliferative, and *Suv4-20h2*-deficient mice are heavier. (A) Samples were collected from unwounded skin and skin 24 hours after incisional wounds were introduced with transmission electron microscopy. The fraction of the nucleus occupied by electron-dense chromatin was plotted. n = 20 nuclei. Mean ± SEM, t-test. Scale bar = 2 µm. **(B)** Excisional 5 mm punch biopsy wounds were introduced into Col1a2-eGFP mice. Samples of skin from unwounded mice, skin far from the wound, and wound-adjacent skin were analyzed with immunofluorescence for CTCF. n = 3. Mean ± SEM, t-test. Scale bar = 50 µm. **(C)** Western blot of H4K20 methylation states in *Suv4-20h2^-/^*^-^ and *Suv4-20h2^+/+^* MEFs. Representative of 3 independent experiments. **(D)** Total cell number in proliferating populations of *Suv4-20h2^-/-^* or *Suv4-20h2^+/+^* MEFs. Two biological replicates. t-tests. **(E)** Fraction of cells in S phase in *Suv4-20h2^-/-^* compared to *Suv4-20h2^+/+^* MEFs. Three biological replicates. t-tests. (F) *Suv4-20h2^-/-^* and *Suv4-20h2^+/+^* MEFs were analyzed with real-time PCR in P and CI conditions for cyclins D2 and E1. Data from three independent MEF lines. Mean ± SEM. t-tests. (G) *Suv4-20h2^-/-^* and *Suv4-20h2^+/+^* MEFs were analyzed with real-time PCR in P and CI conditions for four collagen genes upregulated with CI. Data from three independent MEF lines are shown. Mean ± SEM, t-tests. (H) *Suv4-20h2^+/+^* and *Suv4-20h2^-/-^* mice were weighed weekly from 3 to 7 weeks of age. n = 67-86. Mean ±- SEM, t-tests. **(I)** Weights of *Suv4-20h2^+/+^* and *Suv4-20h2^-/-^* mouse embryos were on day E14-15 are plotted. n = 15-19, mean ± SEM, t-test.

To monitor the levels of CTCF in fibroblasts in a mouse wound healing model, we introduced 5 mm full thickness punch biopsy excisional wounds into the backs of mice genetically engineered to express the ZsGreen fluorescent protein in a fibroblast-specific manner. Mouse skin fibroblasts proximal to wounds contained elevated CTCF levels compared with fibroblasts distant from the wound or fibroblasts from mice that were not wounded (Fig. 7b). Together, these data suggest that quiescence-associated changes in chromatin compaction and CTCF levels in cultured primary human dermal fibroblasts also occur when dermal fibroblasts are activated to proliferate in mouse wounds *in vivo*.

### Suv4-20h2 knockout causes hyperproliferation in mouse embryonic fibroblasts, and results in larger embryos and postnatal mice

Suv4-20h1 and Suv4-20h2 double-knockout mice and Suv4-20h1 knockout mice show developmental delay and die perinatally.^43^ Suv4-20h2-deficient mice are healthy and viable with lower levels of H4K20me3 in several cell populations.^43^ We confirmed *Suv4-20h2*^-/-^ mouse embryonic fibroblasts (MEFs) contain reduced H4K20me3 levels, with little impact on H4K20me2 (Fig. 7c), and absent *Suv4-20h2*, but not *Suv4-20h1*, transcript levels (Supplementary Fig. S8c). *Suv4-20h2^-/-^* MEFs proliferated more rapidly than controls (Fig. 7d), and a higher fraction of *Suv4-20h2^-/-^* MEFs were in S phase in proliferating and contact-inhibited conditions than wild-type controls (Fig. 7e). Transcript levels of cell cycle proteins cyclins D2 and E1 were elevated in *Suv4-20h2^-/-^* compared with *Suv4-20h2^+/+^* MEFs (Fig. 7f). Transcript levels for collagen genes col5a3, col6a6, col9a2, and col23a1 were elevated in contact-inhibited MEFs, and this induction was reduced in contact-inhibited Suv4-20h2 knockout MEFs (Fig. 7g). *Suv4-20h2*-knockout mice were heavier than littermate controls from weeks 3 through 7 of age, consistent with increased proliferation in mice with lower levels of H4K20me3 (Fig. 7h). Further, *Suv4-20h2^-/-^* E14-E15 embryos were heavier than *Suv4-20h2^+/+^*embryos (Fig 7i), consistent with superfluous proliferation in *Suv4-20h2*-deficient embryos.

## Discussion

Our studies reveal a novel mechanism in which H4K20me3 reversibly antagonizes the activity of CTCF to regulate the transition between proliferation and quiescence. Proliferating cells have higher levels of CTCF, lower levels of H4K20me3, more open chromatin, larger, rounder nuclei, larger nucleoli, reduced p27, and increased expression of proliferation-associated genes.

Quiescent fibroblasts, in contrast, have lower levels of CTCF, elevated H4K20me3, more compact chromatin, smaller more elliptical nuclei, smaller nucleoli, and increased expression of extracellular matrix genes. Our data provide evidence that the transition between CTCF and H4K20me3 occupancy at TAD boundaries is a critical contributor to the transition between proliferation and quiescence, and that this switch acts to reprogram gene expression by affecting the functional outcome of existing TADs. Our data further show that in the context of wound healing, nuclear CTCF levels increase in fibroblasts activated to proliferate, and that during embryonic and postnatal development, H4K20me3 restrains organismal growth.

Our findings support a causal and inverse relationship between H4K20me3 and CTCF levels. Knocking down CTCF resulted in a higher fraction of histones with the H4K20me3 modification, while preventing H4K20me3 deposition resulted in more CTCF-bound chromatin, without affecting total CTCF protein abundance. The balance between these two factors was determined by proliferative state, with CTCF higher in proliferating cells and H4K20me3 elevated in quiescent cells. We further show that this CTCF-H4K20me3 axis can affect the proliferation-quiescence decision point, as knockdown of CTCF resulted in slower proliferation and increased levels of quiescence marker p27, while inactivation of an H4K20me3 methyltransferase (Suv4-20h2) resulted in increased proliferation. Strikingly, overexpressing CTCF was sufficient to reduce H4K20me3, promote proliferation in the presence of quiescence signals, and reduce p27 levels.

H4K20me3 has been associated with facultative heterochromatin and chromatin compaction.^7,29,30^ H4K20me3 increases with serum starvation^29^ and silences transcription of genes encoding rRNAs.^29,32,33^ In contrast, we found that genes with promoter-bound H4K20me3 in the quiescent state were evenly split among those that increased and those that decreased in expression when fibroblasts entered quiescence, consistent with an earlier study in which H4K20me3 at promoters did not correlate with gene expression changes with senescence.^44^ Our findings reveal a novel function of H4K20me3 in regulating gene expression by modulating CTCF occupancy at TAD boundaries, consistent with an earlier study that found that H4K20me3 is present at contact domains on chromosome 19 (Ref^45^). Consistent with CTCF’s architectural role,^28^ the factors with the strongest overlap with H4K20me3 peaks were cohesin components SMC1A and SMC3. This finding is also consistent with a previous study that showed Suv4-20h2 knockout embryonic stem cells undergo changes in fine-scale chromatin architecture in gene regulatory regions.^35^

A key finding from our study is the inverse relationship between the abundance of H4K20me3 and CTCF. Our data show that on a chromosome-level scale, large genomic regions can be divided into CTCF-rich, H4K20me3-depleted sections, and distinct CTCF-depleted, H4K20me3-rich portions. The hypothesis that H4K20me3 and CTCF are antagonistic modifiers of chromatin function is further supported by their localization to spatially distinct nuclear territories. Super-resolution microscopy revealed distinct and nonoverlapping nuclear domains occupied by either CTCF or H4K20me3. Our findings are in accord with an earlier demonstration that CTCF and epigenetic marks can display mutual exclusivity.^46^ In our study, H4K20me3-rich regions were compact while the CTCF-rich regions were larger and more diffuse. This may reflect a tendency for H4K20me3-decorated chromatin to self-associate, condense, and separate from domains rich in CTCF. Consistent with our observations, SUV4-20H2 demonstrates phase separations that contribute to compartmentalization of H4K20me3-rich regions.^47^ At a finer scale, during the transition between proliferation and quiescence, thousands of individual loci lose bound CTCF and these same regions, on average, acquire a peak of trimethylated H4K20 centered on the CTCF site.

Several molecular mechanisms could contribute to the H4K20me3-CTCF inverse relationship. DNMT1 can read H4K20me3 and methylate H4K20me3-rich DNA,^48^ and DNA methylation inhibits CTCF binding.^49^ Conversely, TAD establishment is influenced by histone variants^50^ and CTCF can recruit histone demethylases^51^ that may prevent H4K20me3 accumulation at the same genomic loci. Alternatively, SUV420H2 can directly bind and be recruited by the Cohesin complex,^52^ which could contribute to H4K20me3 enrichment at the interfaces of Cohesin:CTCF interactions at TAD boundaries.

Our findings are consistent with earlier studies of CTCF and chromatin in quiescence and senescence. Multiple studies have observed elevated H4K20me3 in quiescent and senescent cells.^7,44^ A previous study also reported CTCF levels are 30% lower in senescent fetal lung fibroblasts than in proliferating fibroblasts,^53^ with an earlier study that observed fewer long-range chromosomal interactions and more short-range interactions in quiescent compared with proliferating fibroblasts.^19^ CTCF binding was observed at the same sites in proliferating and senescent fibroblasts, suggesting that quiescence and senescence may share the characteristic of preserved CTCF-mediated TAD boundaries.^53^ However, in contrast to our findings in quiescent cells, senescent cells harbored many sites with stronger CTCF binding relative to proliferating cells, a feature that may distinguish quiescence from senescence.^53^ Finally, an earlier study observed fewer long-range chromosomal interactions and more short-range interactions in quiescent compared with proliferating fibroblasts.^54^ Senescent cells exhibited a similar but more extreme pattern, with even fewer long-range interactions and even more short-range interactions.^54^

Fibroblasts in tissue can range from thin with flat nuclei in collagen-rich areas to plump with round nuclei in proliferative areas.^55^ We found that human dermal fibroblasts displayed coordinated changes in cell and nuclear size and shape with the transition between proliferation and quiescence, similar to the quiescence-associated morphological transitions observed in T cells^56^ and budding and fission yeast.^57,58^ Surprisingly, we found that CTCF knockdown was sufficient to induce fibroblasts and their nuclei into a thinner and more extended shape. *In vivo*, quiescent skin fibroblasts had low levels of nuclear CTCF that increased substantially when the fibroblasts were activated in response to a wound. Thus, the smaller size and increased ellipticity of quiescent fibroblasts and their nuclei may reflect low levels of CTCF.

Similar changes in cell shape, chromatin conformation, gene expression, and functional properties are observed across species and cell types as cells enter quiescence,^12,59^ but the molecular basis for this commonality has not been understood. Our data show that fibroblasts induced into quiescence by either serum starvation or contact inhibition share common molecular and morphological signatures such as smaller and elliptical nuclei, and reduced nucleoli. Fibroblasts induced into quiescence by either method contained elevated H4K20me3 and reduced CTCF, suggesting the H4K20me3-CTCF axis could be involved in regulating the common phenotypes that occur in response to different quiescence signals.

The model we propose in which H4K20me3 competes with CTCF as cells exit the proliferative cell cycle may be crucially important during early development. H4K20me3 levels are low in embryos and increase during fetal development,^60,61^ suggesting a possible role for H4K20me3-CTCF competition in promoting quiescence during development. Consistent with this prediction, three unrelated human patients with disruptive mutations in one copy of CTCF showed growth impairment.^62^ Additional studies are needed to understand whether dysregulation of the CTCF-H4K20me3 axis contributes to human overgrowth syndromes and disorders associated with inappropriate quiescence-proliferation transitions, and whether competition between architectural and epigenetic factors controls other cell fate transitions.

## ONLINE METHODS

### Fibroblast harvest and culture

To isolate primary human dermal fibroblasts, as previously described in detail, we obtained human skin from newborns via the National Disease Research Interchange. Skin samples were cleaned, sterilized, and partially digested in 0.5% dispase solution. The dermis was separated from the epidermis and the epidermis was removed. The dermis was cut into 2-3 mm pieces and placed in 1000 U/mL collagenase 1a solution at 37°C for 1-2 hours with agitation. Ice-cold Dulbecco’s Modified Eagle Medium (DMEM; Fisher Scientific 11965-118) with 7.5% Fetal Bovine Serum (FBS; Thermo Fisher Scientific, MT35010CV) was added to neutralize the collagenase. The cell suspension was filtered through a 70 µm strainer and centrifuged at 150g for 10 min at 4°C. Cell pellets were resuspended in Complete Growth Media (DMEM containing 8% FBS; 1mM sodium pyruvate; 10 mM HEPES buffer, pH 7.5; 2 mM L-glutamine; and 0.1 mM non-essential amino acids in Modified Eagle Medium) and plated on tissue culture plates. Media was aspirated after 24 hours to remove detached cells and fresh Complete Growth Media was added. After attaching, the fibroblasts had a characteristic spindle-like morphology.

The isolated primary human foreskin fibroblasts were cultured in Growth Media (DMEM containing 10% FBS) and grown at 37°C in 5% CO_2_. Cells were cultured at <80% confluence to maintain log growth. Every 2-3 days, fibroblasts were passaged by incubating them with 0.05% trypsin (Thermo Fisher Scientific 15400-054) for 5 min and replating them at approximately 3 x 10^5^ cells per 10 cm^2^ dish. As needed, cells were cryopreserved in freezing medium (DMEM with 10% FBS and 10% DMSO). Experiments were performed on cells with fewer than 15 passages.

### Quiescence induction

To induce serum-starvation, fibroblasts in Growth Media were passaged, plated at ∼30% confluence, and allowed to attach overnight. The next day, the Growth Media was removed and fibroblasts were washed 3x with room temperature Dulbecco’s Phosphate-Buffered Saline (DPBS; Fisher Scientific, 14190-250) and cultured in Serum Starvation Medium (DMEM containing 0.1% FBS). Medium was replaced every two days with fresh Serum Starvation Media for a total of 7 days. Serum-starved cells were restimulated by passaging them and replating them in Growth Media with full serum (10% FBS).

To induce contact inhibition, proliferating fibroblasts were cultured until they were 100% confluent in Growth Media with media changes every 2 days. Once the fibroblasts were 100% confluent, the cells were maintained for another 7 days with media changes every 2 days. Contact-inhibited fibroblasts were restimulated by passaging them and plating them at proliferative density in Growth Media with 10% serum.

### Phase-contrast light microscopy

Fibroblasts and isolated nuclei were imaged on a Thermo Fisher EVOS™ m5000 imager with phase-contrast. Images were saved and exported into FIJI for downstream analysis.^63^

### Confocal microscopy

Confocal fluorescence microscopy was conducted with a Zeiss LSM 980 imaging system using the ZEN™ (blue edition) software suite. Fluorescent samples were mounted with glass coverslips in Prolong™ glass hardset antifade mounting media (Thermo Scientific P36980). Images were acquired at either 20x or 63x magnification with appropriate excitation and emission filter sets for Alexa Fluor 488, Alexa Fluor 594, Alexa Fluor 647, and Hoescht 33342. Images used for comparative analysis (e.g. siControl vs siCTCF, or proliferating vs serum-starved) were taken at identical magnification, resolution, laser power, detector gain, and scan speed.

### Image analysis

All image analysis was performed on raw image files (.czi) in Fiji.^63^ Confocal images were converted to maximum intensity projections before analysis to remove z-dimensional spatial bias. For nuclear mean fluorescence intensity (MFI) analysis, nuclei were threshholded and segmented based on Hoescht 33342 (Thermo Fisher Scientific H3570) intensity and saved as regions of interest (ROIs). ROIs were analyzed for Hoescht 33342 MFI. For Ki67 image analysis, cells were identified as Ki67 positive if their Ki67 MFI was above a threshold that was held constant across all comparisons in an experiment. For nuclear volumes, Hoescht 33342-labeled samples were imaged on an LSM 880 microscope. 3D reconstructions of individual nuclei were performed in Imaris using Hoescht intensity to demarcate nuclear boundaries. To identify long and short axes of nuclei, nuclei were segmented based on Hoescht staining and bounding ellipses were drawn and analyzed using the FIJI particle analyzer. The nuclear aspect ratio was defined as the ratio of the long to the short axis.

### Incucyte cell proliferation analysis

Cells were seeded in 24-well plates and the number of cells per well was monitored over 72 hours with one hour imaging intervals on an Incucyte SX5 instrument.

### Flow cytometry for nuclear size

Fibroblasts were trypsinized as above and pelleted with centrifugation at 300g for 5 min. Pellets were fixed in 4% paraformaldehyde (Thermo Fisher 50-980) for 15 min at room temperature. Fibroblasts were lysed with 0.3% Triton^TM^ X-100 (Fisher Scientific AC32737-1000) in PBS for 10 min on ice and blocked with 5% bovine serum albumin (BSA; Lee BioSolutions 100-30-250 or Fisher Scientific BP9703100) in PBS for one hour. Fibroblasts were stained with ɑ-Ki67 antibodies conjugated to Alexa Fluor 488 (ThermoFisher 53-5698-82) and analyzed with flow cytometry with the Jonsson Comprehensive Cancer Center Flow Cytometry Core Lab. Nuclei were gated to identify the population with low Ki67 staining and 2N DNA content by Hoescht staining. This population was analyzed for forward scatter to quantify nuclear volume.

### RNA-sequencing

RNA-seq was performed on fibroblast lines generated from three different donors. Whole transcriptome sequencing was performed by the David Geffen School of Medicine Department of Pathology Technology Center for Genomics and Bioinformatics (TCGB). RNA was isolated from cell samples using TRIZOL™ reagent (Thermo Fisher 15596026) according to the manufacturer’s recommendation. RNA was quality checked and quantified using Bioanalyzer (Agilent) RNA Integrity Number (RIN) measurements. Stranded RNA sequencing libraries were prepared with poly-A pulldown. Libraries were sequenced on a NovaSeq X Plus instrument at 2 x 50 bp reads.

### RNA-sequencing analysis

NovaSeq reads were aligned to GENCODE human annotation version 29^64^ (GRCh38) and quantified with Salmon.^65^ Gene-level abundances were summarized by tximport,^66^ and DESeq2^67^ was applied to identify differentially expressed genes (DEGs) between different conditions. Lowly expressed genes were filtered out by requiring a minimum of 10 counts in the number of samples in the smallest condition group. Genes differentially expressed between two conditions with DESeq2 adjusted *p* value less than 0.05 and absolute log_2_(fold change) > 1 were considered DEGs. We considered genes to be differentially expressed between proliferating and quiescent fibroblasts if they were identified as DEGs in both the proliferating vs contact-inhibited and the proliferating vs serum-starved comparisons. Gene Ontology enrichment analysis was performed with clusterProfiler^68^ using all genes that passed the DEseq counts filter as a background. To compare genes regulated by two conditions, we used the hypergeometric test (https://cran.r-project.org/web/packages/SuperExactTest/vignettes/set_html.html). A background list of genes expressed in the scenarios considered was used to test for significant overlap between two lists of differentially expressed genes.

### Immunoblotting

Fibroblasts were trypsinized as above, pelleted with centrifugation at 300g for 5 minutes, and flash-frozen. Frozen cell pellets were incubated in Mammalian Protein Extraction Reagent (MPER; Thermo Fisher Scientific 78501) for 10 min on ice, then centrifuged at 12,000g for 15 min to remove debris. For nuclear extracts, lysates were collected using the standard protocol of the Nuclear and Cytoplasmic Extraction kit (NE-PER; Thermo Fisher PI78834). To isolate chromatin, lysates were collected using the ChromaFLASH Chromatin Extraction Kit (Epigentek P-2001-100).

All protein lysates were quantified with the PIERCE BCA Protein assay (Thermo Fisher 23225). Equal amounts of protein were incubated with NuPAGE LDS sample buffer (Thermo Fisher Scientific NP0008) and NuPage Sample Reducing Agent (Thermo Fisher NP0004) before being heated at 70°C for 10 min and loaded onto polyacrylamide gels for separation and blotting. Pierce^TM^ ECL western blotting substrate (Thermo Fisher 32106) was used to detect horseradish peroxidase-conjugated secondary antibodies and blots were developed onto autoradiography film (Genesee Scientific 30-810).

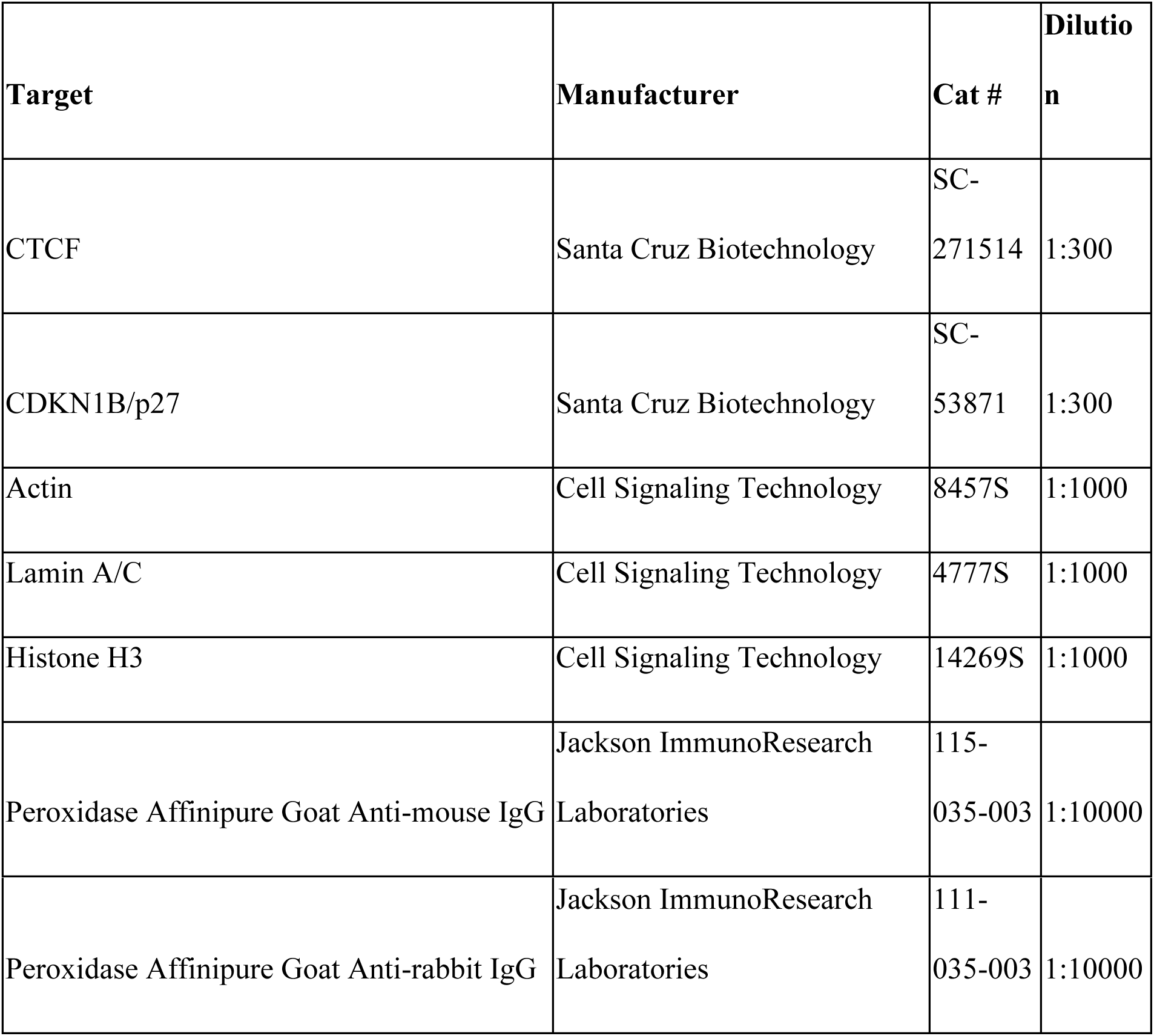

### Immunofluorescence and pyronin Y staining

For immunofluorescence experiments, fibroblasts were plated in 12-well plates with 18mm round coverglass placed at the bottom of each well and coated with gelatin (Sigma Aldrich G2500). Coverslips were collected and fixed in 4% paraformaldehyde (PFA) or ice-cold methanol for 15 minutes (according to antibody specifications) and washed 3x with PBS. Coverslips were permeabilized in 0.3% Triton^TM^ X-100 (Thermo Fisher AC32737-1000) for 15 minutes and then blocked at room temperature for 1 hour in 5% BSA in PBS with 0.2% Tween. Coverslips were incubated with primary antibodies for 1 hour at room temperature or overnight at 4°C followed by 3x wash with PBS. Coverslips were incubated with secondary antibodies for 1 hour at room temperature and washed 3x with PBS. This process was repeated for multiplexed immunofluorescence with primary antibodies from a different host species. Coverslips were treated with 1µg/mL Hoescht 33342 for 15 minutes and washed 3x with PBS before being mounted onto glass slides with Prolong Glass Antifade Mountant (Thermo Fisher P36980). For pyronin Y staining, 2µg/mL of freshly reconstituted pyronin Y (Sigma-Aldrich 83200-5G) was added to the coverslips along with Hoescht 33342 in PBS for 15 min and washed 3x with PBS before mounting.

|  |  |  | **Dilutio** |
| --- | --- | --- | --- |
| **Target** | **Manufacturer** | **Cat #** | **n** |
| H4K20me3 | Active Motif | 91107 | 1:250 |
| CTCF | Cell Signaling | 2899 | 1:250 |
| Fibrillarin | Cell Signaling | 2639S | 1:500 |
| Ki67 | Abcam | ab1667 | 1:500 |
| Goat anti-mouse IgG, alexa fluor 488 | Thermo Fisher | A2817<br>5 | 1:500 |
| Donkey anti-rabbit IgG, Alexa fluor 647 | Thermo Fisher | A3157<br>3 | 1:500 |

### siRNA knockdown

For siCTCF experiments, CTCF siRNA 1 (Sigma Aldrich SASI_Hs01_00230809) and CTCF siRNA 2 (Sigma-Aldrich SASI_Hs02_003 41836), were transfected individually or together into proliferating fibroblasts using Lipofectamine RNAimax™ (Life Technologies 13778-075) following the manufacturer’s recommended protocol. Alternatively, Sigma Aldrich Mission siRNA Universal Negative Control #1 (SIC001-10NMOL) was transfected with Lipofectamine RNAiMax^TM^ according to the manufacturer’s recommended protocol. Briefly, 10 pmol of scrambled or CTCF siRNAs were incubated with Lipofectamine RNAiMax^TM^ reagent in Opti-MEM Reduced Serum Medium (Life Technologies 31985-070) and added dropwise onto fibroblasts plated in a 12-well dish. After cells were incubated in transfection mix for 2 days, the Growth Media was replaced with fresh medium. siRNAs were introduced on day 0 of serum starvation for serum-starved cells.

CTCF siRNAs – #1 CCUAUAUUGAGCAAGGCAA[dt][dt]

#2 CGGUUUACCCAAAGUGGUA[dt][dt]

### Lentiviral overexpression

LTRFORF3290 and TRFORF3550 were gifts from Feng Zhang^69^ obtained through Addgene. Lentiviral particles were generated in accordance with the manufacturer’s guidelines for the ViraPower™ Lentiviral Packaging Mix (Thermo Fisher K497500). 293T cells were plated and transfected with Lipofectamine 2000 (Life Technologies 11668-019) and an optimized mix of 3 packaging plasmids: PLP1, PLP2, and PLP/VSV-G. 2.3µg of PLP1, 1.9µg of PLP2, and 2.7µg of PLP/VSG were mixed with 3µg of lentiviral vector (pLX_TRC317) containing human CTCF ORF (AddGene TFORF3290, plasmid #144766) or mCherry insert (TRFORF3550 Addgene plasmid #145026). Transfection mix was removed 12 hours post-transfection and replaced with fresh Growth Media. Twenty-four to 48 hours after the medium was replaced, viral particles were collected and filtered using a 0.45 µm syringe filter. Fibroblasts were incubated with pools of viral particles for 24 hr. Medium was replaced with medium containing 1µg/mL puromycin (Thermo Fisher A1113803) for selection.

### Super resolution microscopy

Super resolution microscopy was conducted on a Zeiss LSM 980 with Airyscan 2 available through the Broad Stem Cell Center/Molecular Cell and Developmental Biology Imaging Core. Samples were labeled with immunofluorescence and mounted in Prolong™ Glass Antifade Mountant with Neuvitro high-precision coverglass #1 (20mm) (Thermo Fisher NC0308916) to reduce spherical aberrations. Images were acquired with a 63x objective using the Airyscan 2 detector post-calibration and final images were processed using Airyscan processing on the ZEN™ (blue edition) suite.

### CUT&Tag sample preparation and sequencing

CUT&Tag was performed according to the original published protocol^70^ and the manufacturer’s instructions. CUTANA™ CUT&Tag kit was used for H4K20me3 and Active Motif’s CUT&Tag-IT® assay kit with Spike-in control was used for CTCF. In both cases, 500,000 cells were analyzed. Nuclei were isolated and bound to concanvalin beads. Anti-H4K20me3 antibody (1:100 dilution; Abcam AB9053) and anti-CTCF antibody (1:50; Active Motif 61312) were used. H4K20me3 CUT&Tag libraries were generated and sequenced on an Illumina HiSeq X instrument at 2x150bp reads. CTCF libraries were generated and sequenced on an Illumina NovaSeq X Plus instrument at 2x50 bp reads.

### CUT&Tag Analysis

CUT&Tag data was processed following the CUT&Tag Data Processing and Analysis Tutorial^71^ with some modifications. CUT&Tag reads were aligned to the human genome with Bowtie2.^72^ Duplicates were removed with Picard MarkDuplicates.^73^ Mapped reads overlapping with hg38 blacklisted regions^74^ were removed. Peaks were called with SEACR^75^ to identify enriched regions. We selected the top 10% of regions by area under the curve.

Results were analyzed for benchmarks.^75^ We selected a consensus of peaks called in at least 2 samples, across all proliferating, contact-inhibited, and serum-starved replicates. From this consensus, differentially enriched genomic loci were identified using DiffBind^76^ and DESeq2^67^ with adjusted p value < 0.05. We identified 1,149 downregulated and 14 upregulated CTCF peaks. Downregulated CTCF peaks were defined by an absolute log_2_ fold change > 0.5.

Correlations among samples were determined by Pearson correlation of log_2_-transformed read counts. Greater than 25% of peaks overlapped between replicates of H4K20me3 CUT&Tag data in proliferating and of serum-starved fibroblasts. More than 33% of peaks overlapped between replicates of CTCF proliferating, contact-inhibited, and serum-starved fibroblasts. FRiP scores were determined as the fraction of mapped reads in called peaks.

Trackplots were generated by the TRACKPLOT^TM^ Open Source Software. GIGGLE scores were determined with Cistrome.^77^ GIGGLE ranks query results by a composite of the product of −log_10_(p value) and log_2_(odds ratio) to generate a GIGGLE score.^36^

### RNA isolation and real-time-quantitative PCR

RNA was isolated from mouse embryonic fibroblasts using Directzol RNA miniprep kit (Zymo). Following total RNA isolation, 50ng of RNA was used for rt-qPCR using SuperScript III One-Step RT-PCR System (ThermoFisher) following standard protocols. The real-time reaction was performed on a CFX96 Real-Time PCR system (Bio-Rad).

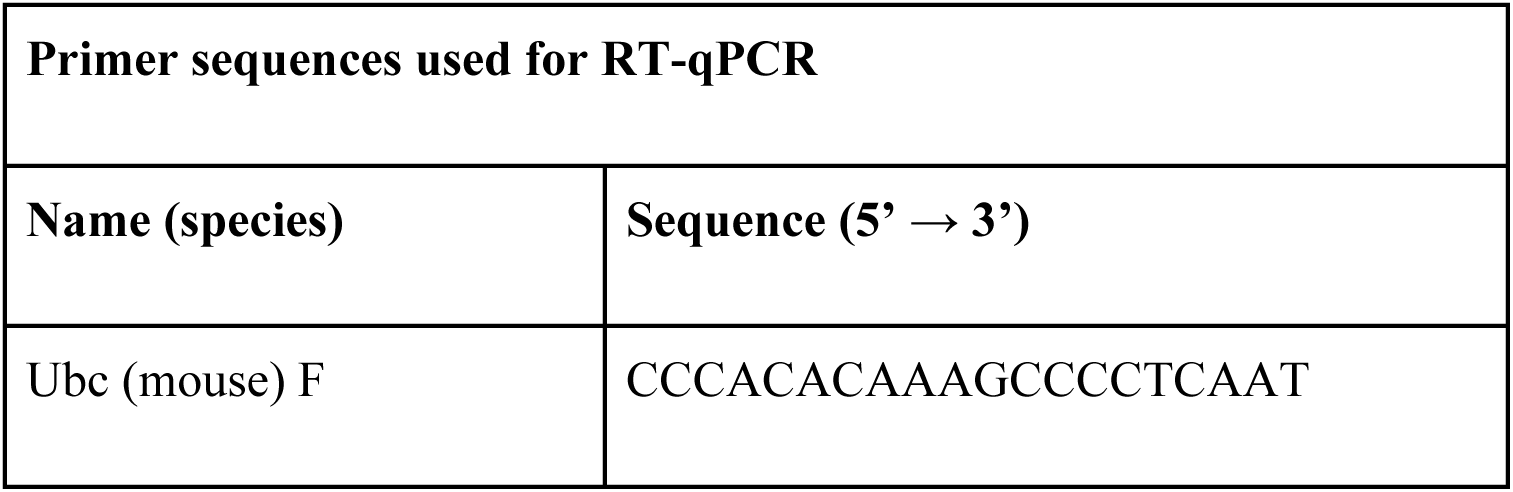
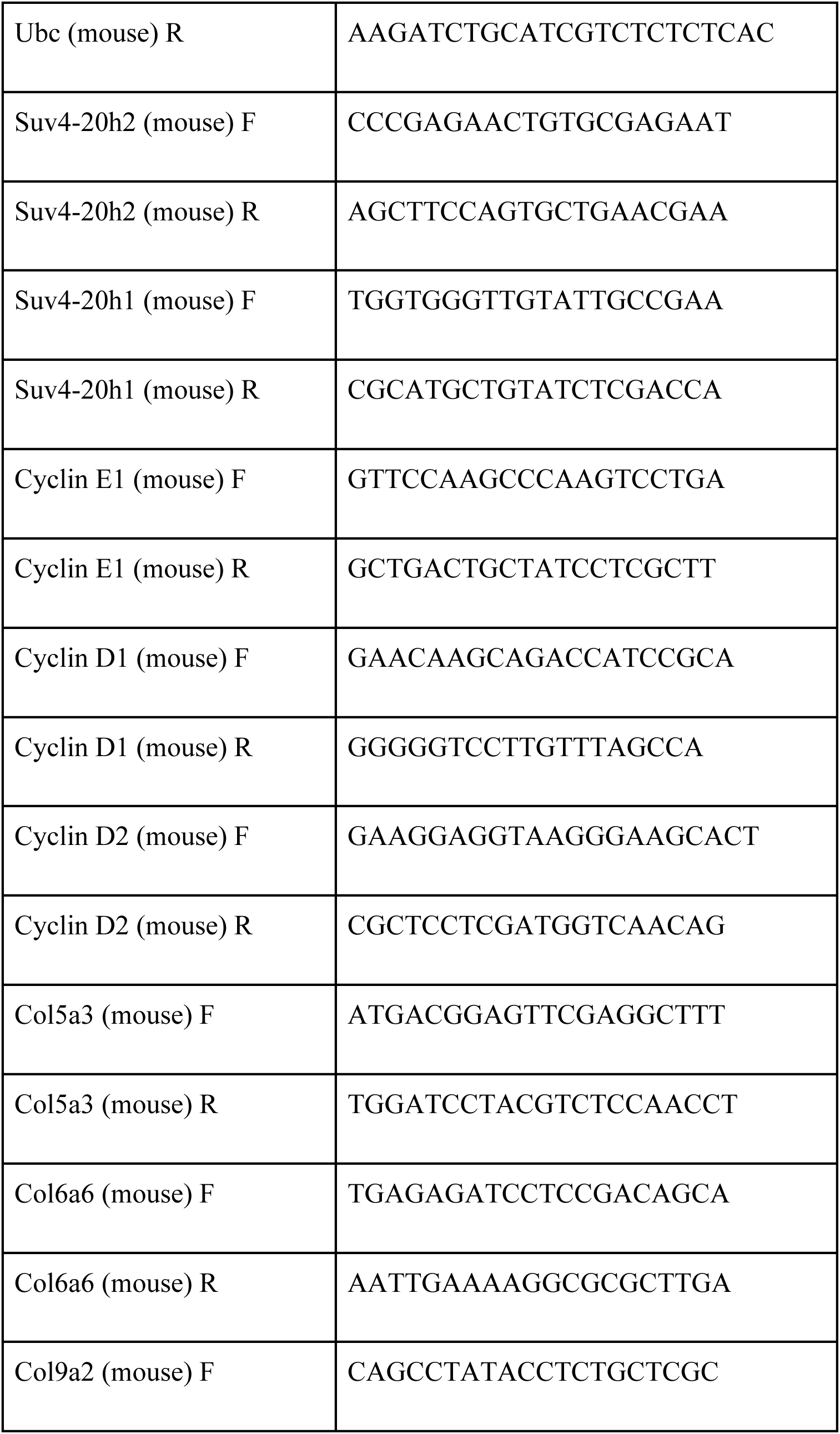
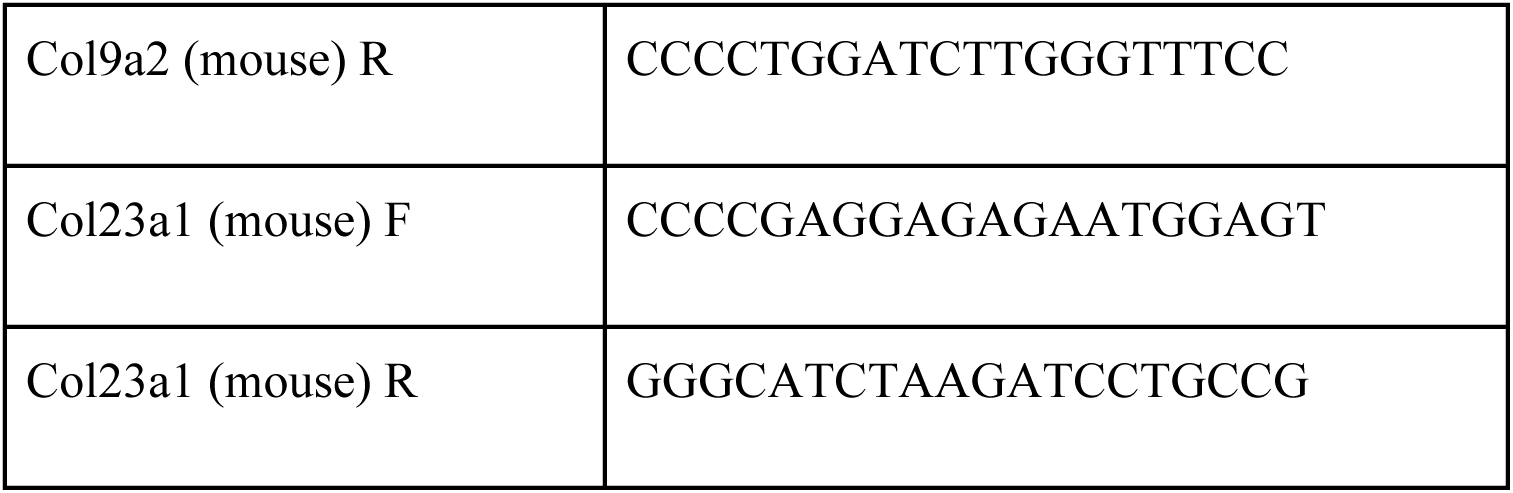

### MEME/SEA

SEACR^75^ was used to identify the top 10% of H4K20me3 peaks in serum-starved fibroblasts. SEA (Simple Enrichment Analysis) was performed on these peaks for the JASPAR CTCF Motif matrix (MA0139.1).^78^

### CTCF-H4K20me3 overlap and proximity to TADs

To assess whether sites with reduced CTCF binding are more likely to be found within 100 kb of a TAD boundary, we considered CTCF binding sites present in at least 2 replicates across all three conditions. To determine the fraction of CTCF and H4K20me3 peaks that overlap, we considered loci with both CTCF and H4K20me3 peaks in at least one of the conditions tested. TAD consensus estimates were downloaded from TAD Map.^37^ H4K20me3 and CTCF CUT&Tag reads within 50 kb were considered close to a TAD boundary. We tested whether TADs demarcated by downregulated CTCF sites were enriched for genes differentially expressed between proliferating and quiescent fibroblasts. To account for variations in gene density between TADs, we tested enrichment by normalizing the number of DEGs by the total number of genes, excluding lowly-expressed genes (see RNA-seq analysis), in each TAD. Differences were assessed with a Wilcoxon rank-sum test for the proportion of genes within demarcated TADs that are DEGs.

### Mouse husbandry

All procedures for incisional wounds were performed in accordance with protocols approved by the Princeton University and University of California Los Angeles Institutional Animal Care and Use Committees. C57BL/6J mice aged 4-5 weeks were used for incisional wounding experiments. Incisional, full thickness wounds were introduced into the dermal skin on the mouse’s back. Col1a2-CreER mice were provided to the Coller laboratory as a generous gift from Dr. Arjun Deb (UCLA) and Dr. Andrew Leask (University of Toronto).^79^ Excisional, full thickness, 0.5 mm punch biopsies were introduced into the dermal skin on the backs of male Col1a2-CreER.ZsGreen mice aged 8-10 weeks. At approximately 6 weeks of age, the mice were treated with tamoxifen *i.p.* (200 µl of 10 mg/ml of corn oil) daily for five days to activate cre recombinase. Suv4-20h2^-/-^ mice^43^ were generously provided by Peter Adams, UCSD, with the approval of Gunnar Schotta.

### Mouse embryonic fibroblast generation

Primary mouse embryonic fibroblast cells were established by standard procedures from *Suv4-20h2^-/-^* and *Suv4-20h2^+/+^* embryos at E13.5 as previously described.^80^ Cells were grown in DMEM (Life Technologies) with 10% fetal bovine serum (Life Technologies) at 5% CO_2_. Contact-inhibited mouse embryonic fibroblasts were grown to full confluency and incubated for 7 to 10 days with medium changes every 3 days.

### Weight experiments

For embryo weights, embryonic day 14-15 embryos were collected and removed from their placentas. Embryos were weighed individually without their placentas. For whole body weights, mice were weaned from their mothers and weights were recorded at weekly post-birth.

### Transmission electron microscopy

Skin samples were fixed in 3% glutaraldehyde in 100mM sodium cacodylate buffer (pH 7.2) and post-fixed in 2% osmium tetroxide for 1 hr on ice. *En bloc* staining with 1% uranyl acetate in 0.05 M sodium maleate buffer was performed for 16 hr at room temperature. Samples were dehydrated with ethyl alcohol and embedded in Epon resin (Electron Microscopy Sciences 14120). For skin sections, the appropriate location within the tissue block was identified by staining 1mm thick sections with 1% Toluidine Blue in water (Electron Microscopy Sciences 22050) to visualize nuclei. 70nm ultra-thin sections were contrasted with lead citrate and imaged on a Zeiss Leo 912AB transmission electron microscope at 80 kV equipped with EMCCD (Hamamatsu Orca I, 2848 x 2626 pixels) and image software (AMT ver. 602).

### Statistics

Bar graphs were plotted as mean ± SEM. For comparison between two groups (*e.g.* siControl vs. siCTCF), two-tailed, unpaired t-tests were used. For comparisons involving multiple samples (*e.g.* proliferating, serum-starved, contact-inhibited samples), one-way ANOVA was performed followed by Tukey’s Honestly Significant Difference (HSD) test. *p*-values less than 0.05 were considered significant. One asterisk indicated p<0.05, 2 asterisks indicated p < 0.01, 3 asterisks indicated p <0.001, 4 asterisks indicated p< 0.0001.

## Data Availability

### CUT&Tag: Zenodo link

https://zenodo.org/uploads/19909506?token=eyJhbGciOiJIUzUxMiJ9.eyJpZCI6ImJlY2ZlMjdlLWJmNDUtNDNiNi1iYjI3LTA5OTk5NDg3Njc4YSIsImRhdGEiOnt9LCJyYW5kb20iOiJiMmJhYWY3YzI4MDBlMzgwNmFiNzIwNGVjOWYzMDNhNyJ9.J3l73k_CdcWfYSpfxirMMmSZDKJU0b_riAJ1CZqdVkyoG9eb-9p-Ls09LvCpjJCViQe7-ALh1wLW5coaeVyRwA

### RNA-seq (triplicates): Zenodo link

https://zenodo.org/uploads/19910556?token=eyJhbGciOiJIUzUxMiJ9.eyJpZCI6ImM0MGZiY2IxLWVlM2YtNGQyMy1iODJhLThlNWNiMWY2MDc2MSIsImRhdGEiOnt9LCJyYW5kb20iOiJmZGZlMjA3ZTUwNDMwY2RlNDIwYjIxYTBkN2M5YWVhNyJ9.k6C8yl3p-MbJohqxVYN8dB7YMdIWTP0am-RFUPWOTi_q_ZTvQcU4lMpy5DE9RKik9W0A4vyFJTgPx5RDPeDUAg

### RNA-seq perturbation (A196, DMSO): Zenodo link

https://zenodo.org/uploads/19909021?token=eyJhbGciOiJIUzUxMiJ9.eyJpZCI6ImYwODM3MDM5LWFkMzUtNDcxMi1iODQyLWNjNWRhMThkMmJmNiIsImRhdGEiOnt9LCJyYW5kb20iOiJmNTdlZDBiOGIwMDQ4ODFkYTU5MTE0MTc0YjJiNDk0YyJ9.Cfq6pqS0X-W6_TKGUFEKa2J-LgSRyTUycuuATHgaud17Z0vop_x3dj8u7vRiR3f_xOwUm_BCmiW2vSnO1VPfYQ

### RNA-seq perturbation (CTCF OE, RFP OE): Zenodo link

https://zenodo.org/uploads/19955121?token=eyJhbGciOiJIUzUxMiJ9.eyJpZCI6ImJiNjVhNTFkLTNjMGQtNGIzOS1hZmUwLTlmYzFiNmVhNzg5YiIsImRhdGEiOnt9LCJyYW5kb20iOiJiNDkyOTI5OGUxZDA4MjEyODYxZWRjMGU4NDUxOTJjYiJ9.GyieHqQCbzul41qSZLqocSS-EIysRW_b8GFIjM2AnWmXdvHiOMYd0CCIayRSkv1A3M9Je32X_oW6eAeFuVBBTg

### RNA-seq perturbation (siCTCF, siControl): Zenodo link

https://zenodo.org/uploads/19955327?token=eyJhbGciOiJIUzUxMiJ9.eyJpZCI6ImFkM2U4MjVkLTc2YzctNGQ5Yi05MjA0LTcwOGQzOGRkOTU4NyIsImRhdGEiOnt9LCJyYW5kb20iOiI5YTU5NmQyYTA3NzVhMDFmNjhhYmEyYjE3MWQ2NWRlOSJ9.79HjyWxsGz1b1saqxOajL9wT-sPKSEEG4TlyQkf7xO8zQOL6mPxHME8yZk3fdXH2Pp0GVWSy6s8170XYLK9BYg

### Code repository: github link

https://github.com/20richardh/collerlab-h4k20me3-ctcf-rnaseq-cutandtag.git

## Acknowledgements

This work was supported by grants to HC NIGMS R01 GM081686, NIGMS R01 GM0866465, NIH R01 AR070245, NCI RC1 CA147961, 1R01 AR084245, the Cancer Research Institute CLIP grant, a Melanoma Research Alliance Team Science Award, a Melanoma Research Foundation Award, the Iris Cantor Women’s Health Center/UCLA CTSI NIH Grant UL1TR000124, the UCLA SPORE in Prostate Cancer (P50 CA092131), David Geffen School of Medicine Metabolism Theme, University of California Cancer Research Coordinating Committee, Broad Stem Cell Center Innovation Awards, and Rose Hills Foundation and Hal Gaba awards from the UCLA Broad Stem Cell Center, a Jonsson Comprehensive Cancer Center Seed Grant and Leader’s Vision Awards. KA acknowledges support through the UCLA Molecular Biology Institute Whitcome Fellowship and UCLA Center for Duchenne Muscular Dystrophy Ruth L. Kirschstein National Research Service Award T32AR065972 Muscle Cell Biology, Pathophysiology, and Therapeutics from the National Institute of Arthritis and Musculoskeletal and Skin Diseases. This work was funded by the Cell and Molecular Biology Training Grant (NIH Ruth L. Kirschstein National Research Service Award GM007185; to A.Z.C.). AZC acknowledges a UCLA Molecular Biology Institute Whitcome Fellowship. AA acknowledges support from the UCLA Dermatology T32 (T32AR071307), the UCLA Tumor Cell Biology T32 (5T32CA009056-038), and a Broad Stem Cell Center Postdoctoral Fellowship in Stem Cell Science. HAC was the Milton E. Cassel scholar of the Rita Allen Foundation.

## Author contributions

K.A., M.M., E.B., R.H., A.C., D.J., L.V., A.A., K.S., W.M.S., K.B., A.D, and H.A.C contributed to the conception and design of the work. K.A., M.M., E.B., R.H., A.C., D.J., L.V., A.A., K.S., W.M.S., K.B., A.D, and H.A.C contributed to the acquisition, analysis, or interpretation of data. M.M, R.H., and W.J.M. wrote computer code to analyze data. K.A., M.M., E.B., R.H., A.C., D.J., L.V., A.A., K.B., and H.A.C. contributed to drafting and revising the work. All authors approved the submitted version.

## Ethics declaration and consent to participate

Discard human skin collection was approved by the Princeton University Institutional Review Board (protocol 3134) and the UCLA Institutional Review Board (14-0145). Informed consent was obtained by the National Disease Research Interchange. Animal experimentation was approved by the Princeton University Office for Animal Research (1813) and the UCLA Office for Animal Research (2014-049, 2015-033). Mice were euthanized in accordance with institutional and ethical guidelines. All experimental methods comply with the Helsinki Declaration.

## List of Supplementary Tables

**Table S1 Quantified transcript data from RNA-seq proliferating versus quiescence**

**Table S2 Genes significantly differentially expressed in proliferating versus quiescent fibroblasts**

**Table S3 Quantified transcript data for genes in the GO category chromatin remodeling**

**Table S4 Quantified transcript data from CUT&Tag fastq/bam files, SEACR peaks**

**Table S5 DiffBind results from CUT&Tag data**

**Table S6 Quantified transcript data from RNA-seq from siCTCF, CTCF overexpression and A-196 experiment**

**Table S7 Genes significantly differentially expressed in siCTCF, CTCF overexpression and A-196**

## Supplementary Figures

**Supplementary Figure S1.**
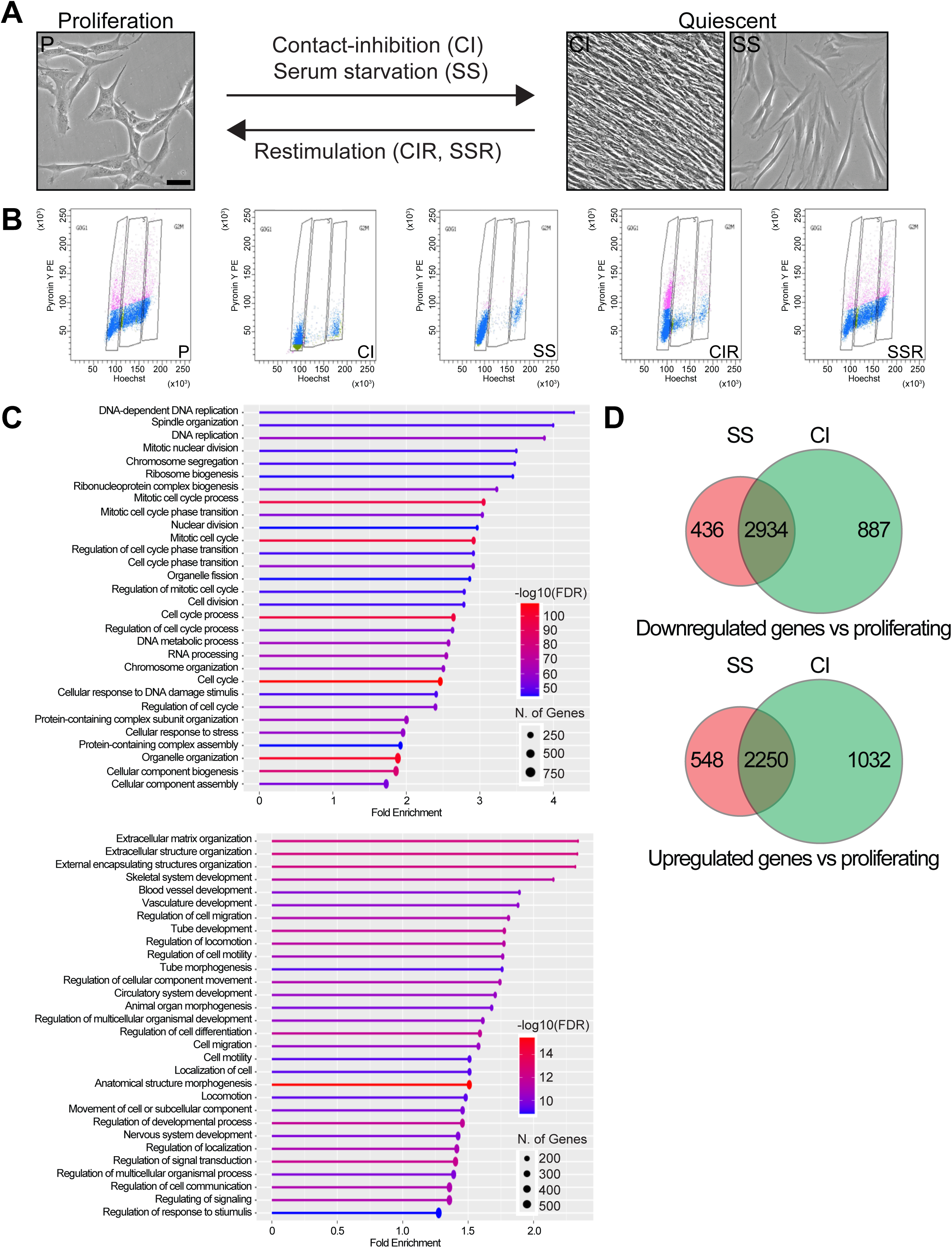
Gene Ontology enrichment for quiescent cells reveals downregulation of cell cycle genes, upregulation of extracellular matrix genes, and substantial overlap between genes regulated in serum starvation and contact inhibition conditions. Related to Figure 1. **(A)** Primary human dermal fibroblasts were imaged with light microscopy in proliferating conditions and quiescent conditions induced by serum starvation or contact inhibition. **(B)** Proliferating (P), serum-starved (SS), contact-inhibited (CI), serum-starved and restimulated (SSR), and contact-inhibited and restimulated (CIR) fibroblasts were stained with pyronin Y (y-axis) and Hoechst 33342 (x-axis) and analyzed with flow cytometry. **(C)** RNA-seq was performed on three independent strains of proliferating (P), serum-starved (SS), contact-inhibited (CI), serum-starved and restimulated (SSR), and contact-inhibited and restimulated (CIR) fibroblasts. Differentially expressed genes in both serum-starved and contact-inhibited fibroblasts (quiescent fibroblasts) compared with proliferating fibroblasts were identified. Gene Ontology terms enriched in genes downregulated (top) and upregulated (bottom) in quiescent compared with proliferating fibroblasts are shown. **(D)** Venn Diagram showing the overlap among Differentially expressed genes downregulated or upregulated in serum-starved compared with proliferating fibroblasts and Differentially expressed genes downregulated in contact-inhibited compared with proliferating fibroblasts.

**Supplementary Figure S2.**
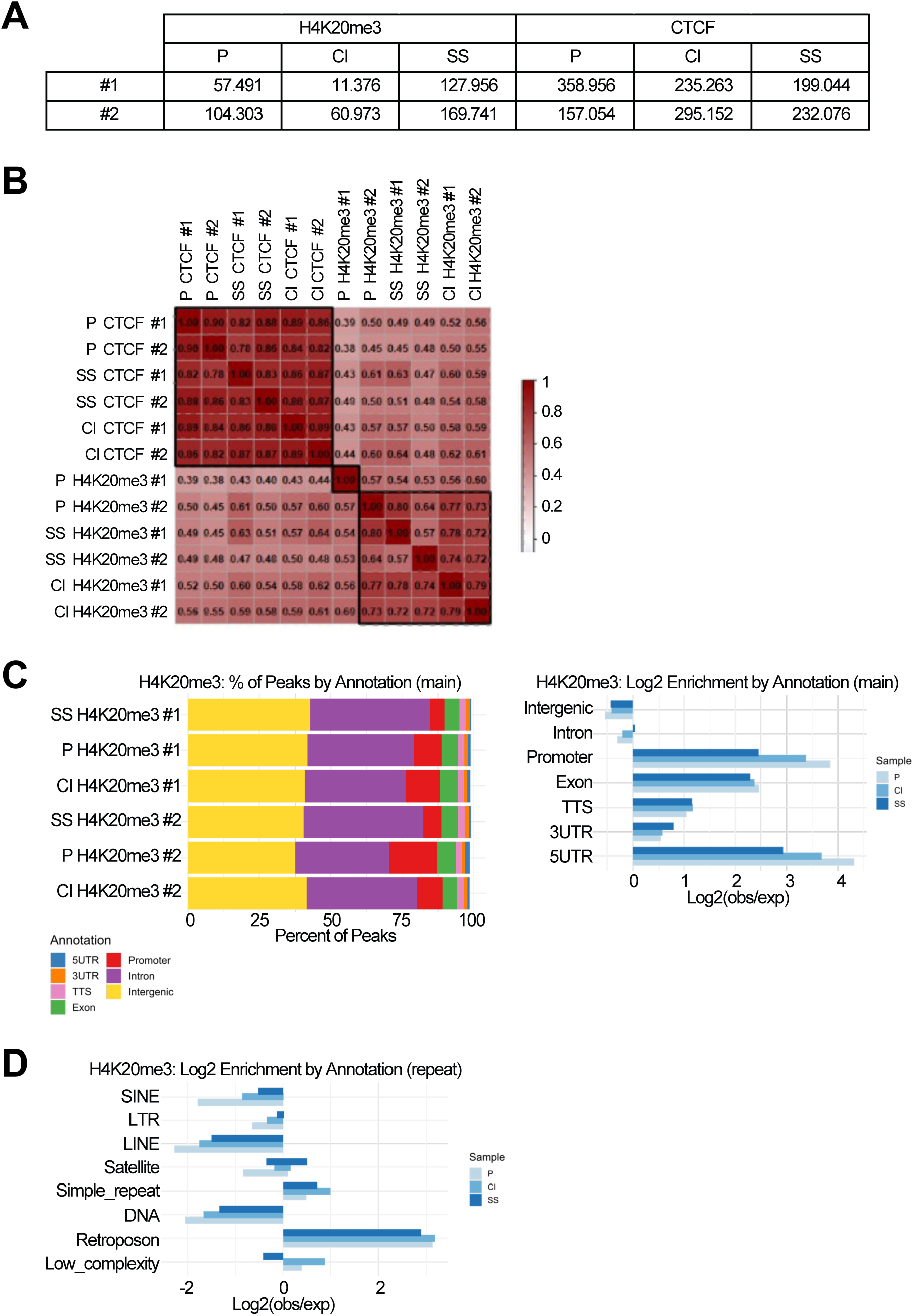
Genomewide distribution of H4K20me3 in proliferating and quiescent fibroblasts. Related to Fig. 1. **(A)** CUT&Tag was performed on proliferating, serum-starved, and contact-inhibited fibroblasts derived from two different donors for H4K20me3 and CTCF. The numbers of peaks in each sample for each condition are provided. **(B)** Pearson correlations were performed among every pairwise comparison of CUT&Tag samples. Samples were hierarchically clustered. Each comparison (row and column) is colored based on the extent of correlation of the peaks according to the scale bar shown with blue indicating anti-correlation and red indicating a positive correlation. **(C)** H4K20me3 ATAC-seq peaks were mapped to the human genome assembly hg38. The distribution of peaks among different types of regions was plotted (left). Log base_2_ fold-enrichment (observed/expected) for different genomic regions is shown for proliferating (P), serum starved (SS), and contact inhibited (CI) fibroblasts (right). **(D)** H4K20me3 ATAC-seq peaks were associated with a RepeatMasker bed file of mapped repetitive regions in hg38. Log base_2_ fold-enrichment for H4K20me3 ATAC-seq peaks in different types of repeats is shown for proliferating (P), serum starved (SS) and contact inhibited (CI) fibroblasts.

**Supplementary Figure S3.**
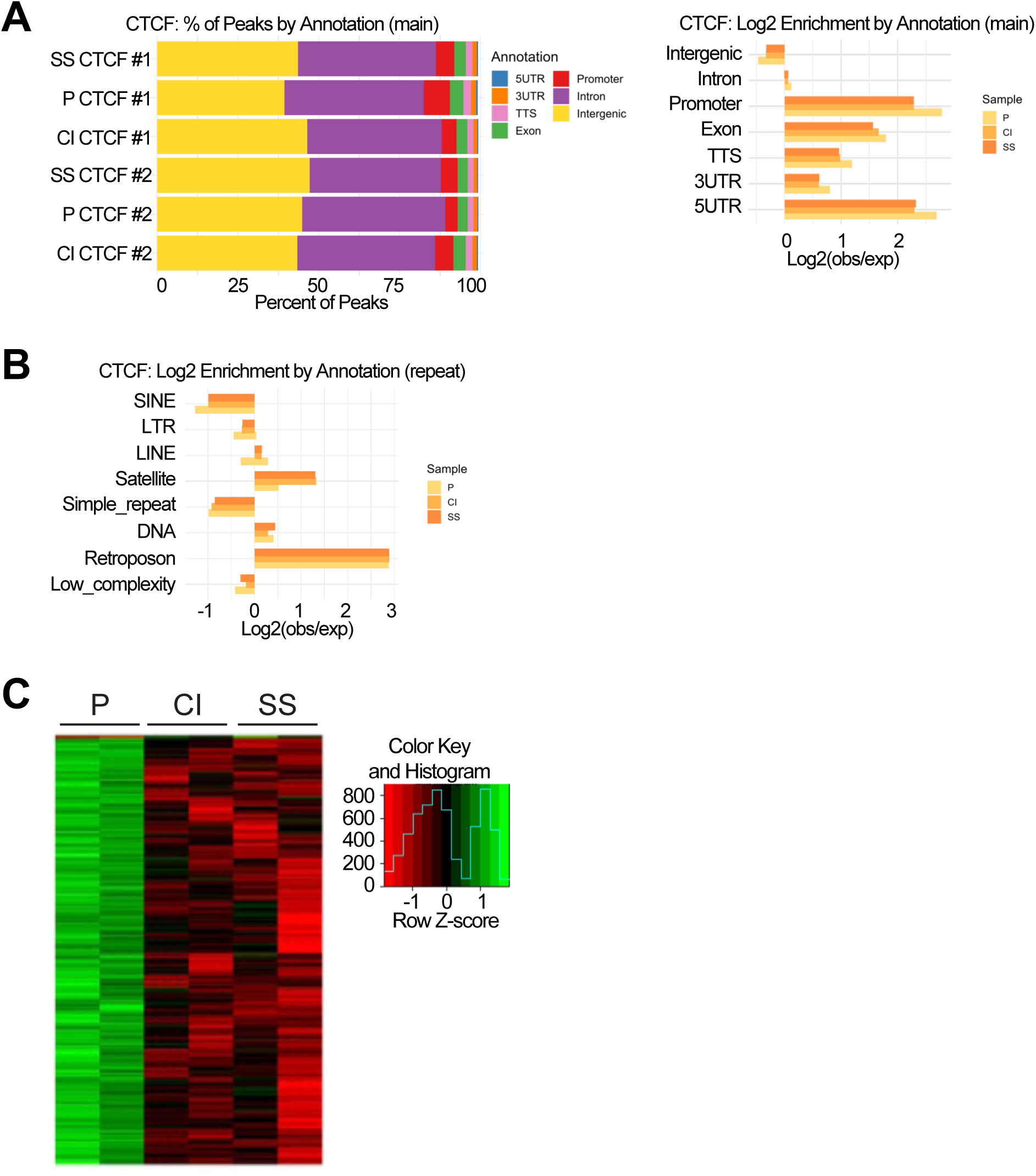
Genomic distribution of CTCF-bound chromatin in proliferating and quiescent fibroblasts and decreased CTCF binding in quiescent than proliferating fibroblasts. Related to Fig. 2. **(A)** CTCF ATAC-seq peaks were mapped to human genome assembly hg38. The distribution of peaks among different types of regions was plotted for two strains of fibroblasts in proliferating (P), serum starved (SS) and contact inhibited (CI) conditions (left). Log base_2_ fold-enrichment (observed/expected) for different genomic regions is shown for proliferating (P), serum starved (SS) and contact inhibited (CI) fibroblasts (right). **(B)** The locations of CTCF ATAC-seq peaks were associated with a RepeatMasker bed file of mapped repetitive regions in hg38. Log base_2_fold-enrichment (observed/expected) for CTCF ATAC-seq peaks in different types of repeats in proliferating (P), serum starved (SS) and contact inhibited (CI) fibroblasts are plotted. **(C)** CTCF CUT&Tag data for two replicates each of proliferating (P), serum-starved (SS) and contract-inhibited (CI) fibroblasts are presented in heat map format. Normalized peak intensity for each sample is indicated by color based on the color scale provided.

**Supplementary Figure S4.**
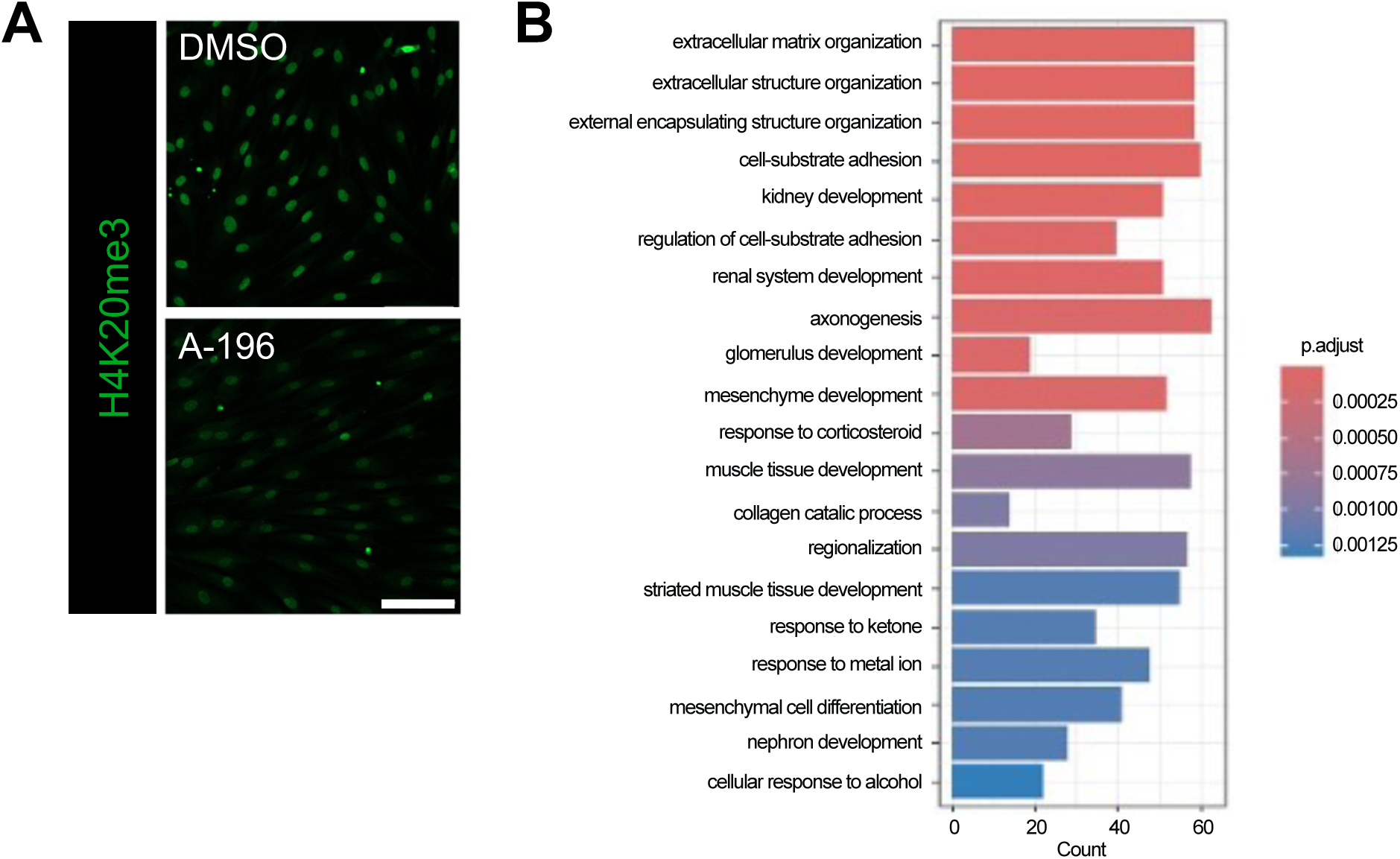
A-196 treatment reduces H4K20me3 and results in downregulation of extracellular matrix-related genes. Related to Figure 3. **(A)** Serum-starved fibroblasts were treated with A196 or DMSO as vehicle control. H4K20me3 was monitored with immunofluorescence. Quantification is provided in Fig 2a. Scale bar = 100 µm. **(B)** Differentially expressed genes that are upregulated in serum-starved fibroblasts compared with proliferating fibroblasts and downregulated in fibroblasts treated with A-196 versus DMSO were identified. Gene Ontology categories enriched in these genes are shown. The number of genes is plotted on the x-axis and the statistical significance is shown by the color according to the color scale.

**Supplementary Figure S5.**
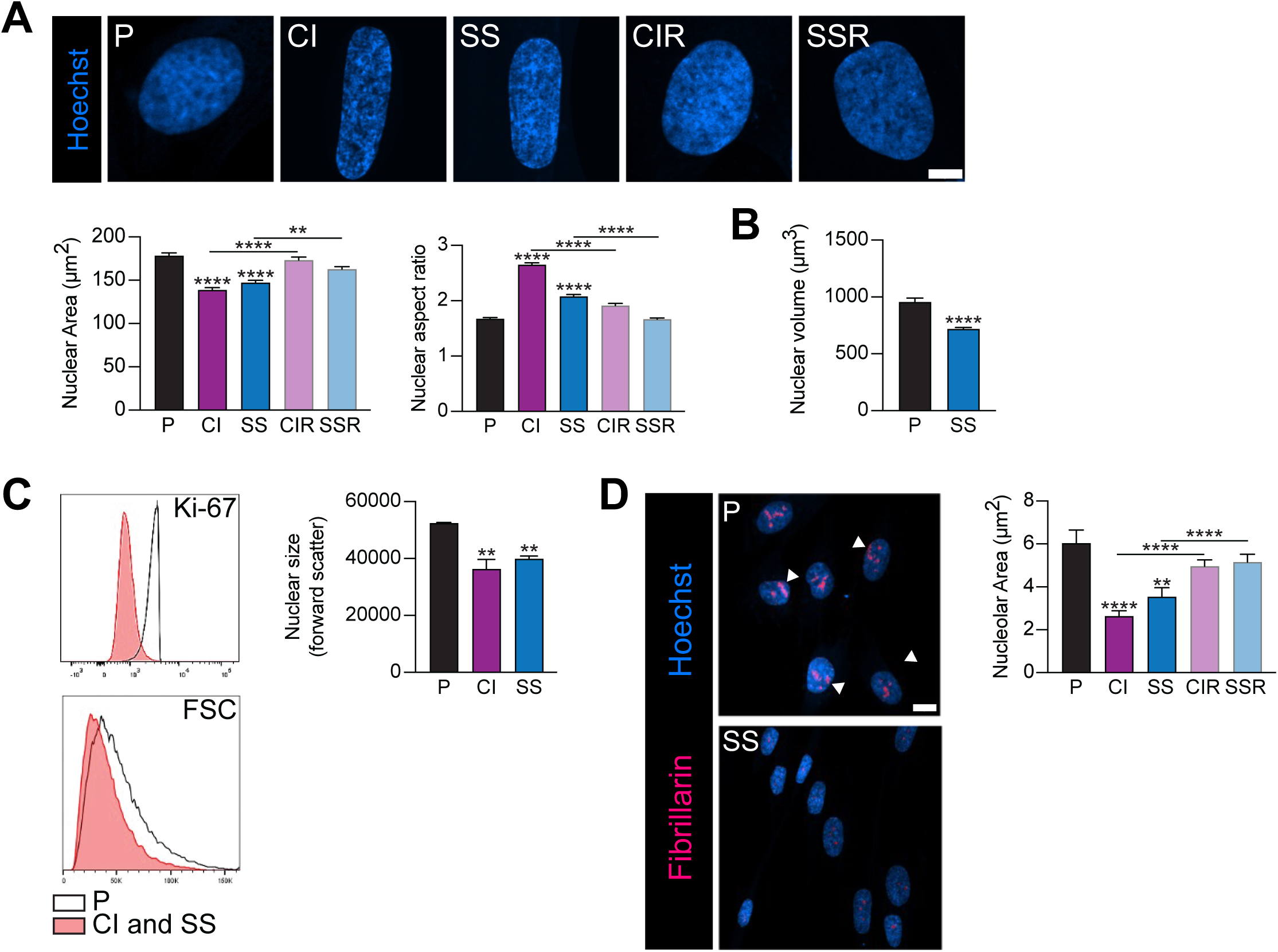
Fibroblast nuclei are smaller and elliptical, and nucleoli are smaller when fibroblasts reversibly enter quiescence by serum starvation or contact inhibition. Related to Figure 3. **(A)** Fibroblasts were stained with Hoescht and imaged with confocal microscopy in proliferating (P), serum-starved (SS), contact inhibited (CI), serum-starved and restimulated (SSR), and contact-inhibited and restimulated (CIR) states. n = 200-400. Mean +/- SEM, ANOVA. Scale bar = 5 µm. **(B)** Proliferating and serum-starved fibroblasts were stained with Hoescht and rhodamine phalloidin and imaged. Three-dimensional nuclei were generated with Imaris. Nuclear volumes in proliferating (P) and serum-starved (SS) fibroblasts were plotted. n = 70-110. Mean +/- SEM, t-test. **(C)** Nuclei liberated from proliferating (P), contact inhibited (CI), and serum-starved (SS) fibroblasts were stained with Hoescht for DNA content and a fluorescent antibody against Ki-67 and analyzed with flow cytometry. A histogram of Ki-67 levels in proliferating (P), and a combination of serum-starved and contact-inhibited populations is shown (left, top). n = 10,000 events in duplicate samples. A histogram of forward scatter values for Ki67-negative fibroblasts with 2N DNA content from proliferating and quiescent cells is shown (left, bottom). n = 10,000 events in duplicate samples. Nuclear size as determined by flow cytometry is plotted for proliferating (P), serum-starved (SS), and contact-inhibited fibroblasts (CI) (right). n = 10,000 events in duplicate samples. Mean +/- SEM, ANOVA. **(D)** Proliferating (P), contact-inhibited (CI), serum-starved (SS), contact inhibited restimulated (CIR), and serum starved restimulated (SSR) fibroblasts were analyzed with immunofluorescence for fibrillarin to mark nucleoli. Nucleolar area (µm^2^) was plotted. n = 100-600 cells. Mean +/- SEM, ANOVA. Scale bar = 10 µm.

**Supplementary Figure S6.**
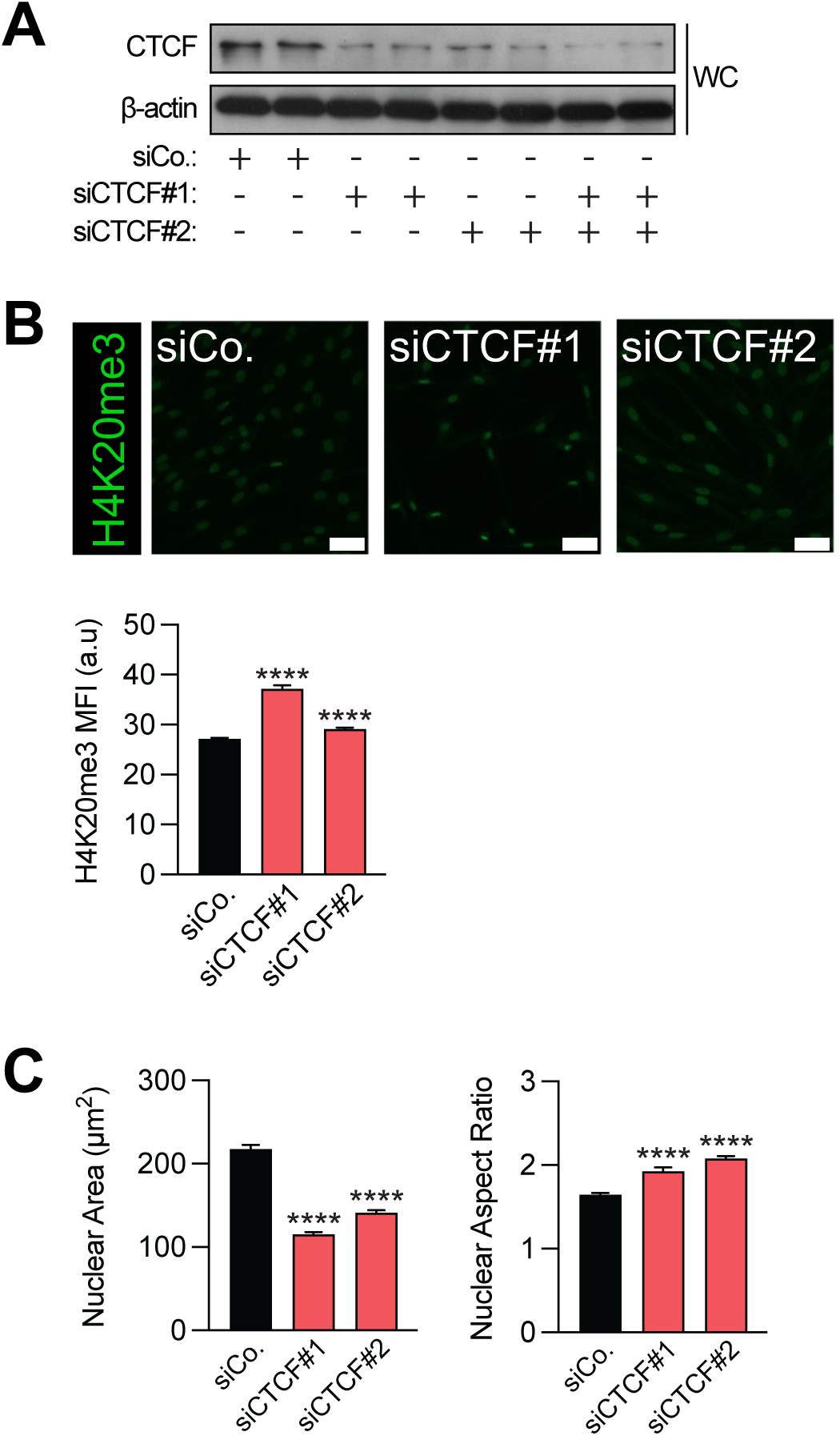
Each of two siRNAs against CTCF produce effective knockdown, increased H4K20me3, reduced nuclear area, and an increased nuclear aspect ratio. Related to Figure 4. **(A)** Fibroblasts were transfected in duplicate with a control siRNA, siRNA to CTCF #1, siRNA to CTCF #2, or a combination of siRNAs to CTCF #1 and #2. Two days later, protein lysates were isolated and CTCF levels were determined with immunoblotting. A portion of this western blot is shown in Fig. 4A. **(B)** Proliferating fibroblasts were transfected with a control siRNA, siRNA to CTCF #1, or siRNA to CTCF #2. Fibroblasts were analyzed with immunofluorescence for H4K20me3. n = 140-400 cells. Mean +/- SEM, ANOVA. Scale bar = 50 µm. **(C)** Proliferating fibroblasts were transfected with a control siRNA, siRNA to CTCF #1, or siRNA to CTCF #2. Nuclei were stained with Hoescht and imaged with confocal microscopy. n = 140-400 cells. Mean +/- SEM, ANOVA.

**Supplementary Figure S7.**
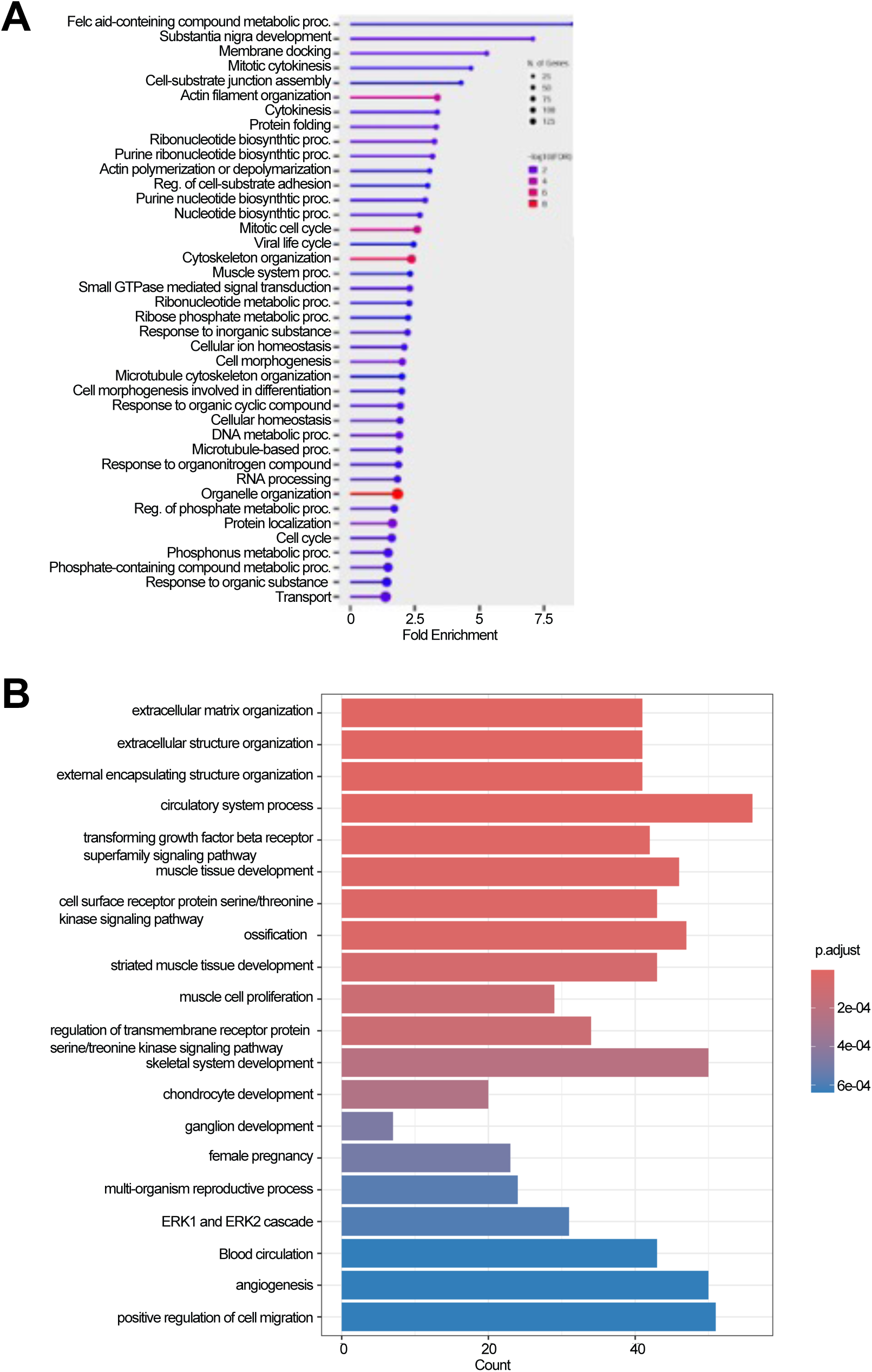
Gene Ontology categories for genes regulated with quiescence and CTCF knockdown are enriched for nucleotide synthesis and extracellular matrix production. Related to Figure 4. **(A)** Differentially expressed genes that are downregulated in serum-starved compared with proliferating fibroblasts and downregulated in fibroblasts transfected with siRNAs against CTCF versus a control siRNA were identified. Gene Ontology categories enriched in these genes are shown. Fold-enrichment is indicated in the x axis. The number of genes is indicated by the size of the circle. Statistical significance is shown by the color as in the color scheme. **(B)** Differentially expressed genes that are upregulated in serum-starved fibroblasts compared with proliferating fibroblasts and upregulated in fibroblasts transfected with siRNAs against CTCF versus a control siRNA were identified. Gene Ontology categories enriched in these genes are shown. The x axis indicates the number of genes. Statistical significance is indicated by the color as shown in the color scheme.

**Supplementary Figure S8.**
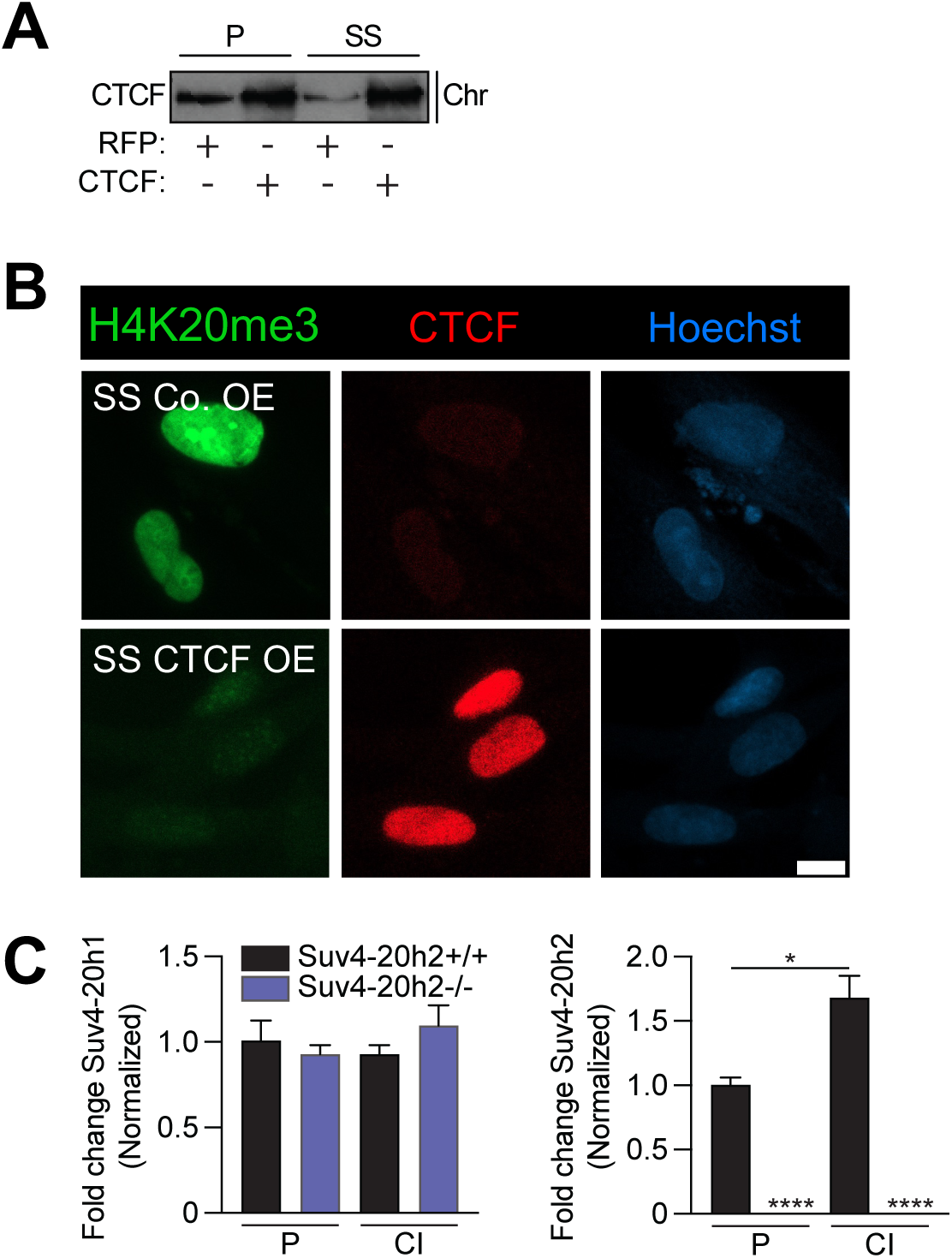
Validation of CTCF overexpression and Suv4-20h2 knockout MEFs. Related to Figure 5. **(A)** Serum-starved fibroblasts were transduced with a lentiviral vector overexpressing CTCF or a control protein and analyzed with immunoblotting for CTCF. **(B)** Serum-starved fibroblasts were transduced with a lentiviral vector overexpressing CTCF or a control protein and analyzed with immunofluorescence for H4K20me3 and CTCF. **(C)** RNA isolated from mouse embryonic fibroblasts generated from *Suv4-20h2^-/-^* (*Suv/suv*) or *Suv4-20h2^+/+^* mice was analyzed with real-time PCR for *Suv4-20h1* (left) and *Suv4-20h2* (right) transcript levels.

## References

1. Dixon, J. R. et al. Topological domains in mammalian genomes identified by analysis of chromatin interactions. Nature 485, 376–380 (2012).

2. Symmons, O. et al. Functional and topological characteristics of mammalian regulatory domains. Genome Res. 24, 390–400 (2014).

3. Kim, T. H. et al. Analysis of the Vertebrate Insulator Protein CTCF-Binding Sites in the Human Genome. Cell 128, 1231–1245 (2007).

4. Smith, E. M., Lajoie, B. R., Jain, G. & Dekker, J. Invariant TAD Boundaries Constrain Cell-Type-Specific Looping Interactions between Promoters and Distal Elements around the *CFTR* Locus. Am. J. Hum. Genet. 98, 185–201 (2016).

5. Takayama, N. et al. The Transition from Quiescent to Activated States in Human Hematopoietic Stem Cells Is Governed by Dynamic 3D Genome Reorganization. Cell Stem Cell 28, 488–501.e10 (2021).

6. Kallingappa, P. K. et al. Quiescence Loosens Epigenetic Constraints in Bovine Somatic Cells and Improves Their Reprogramming into Totipotency1. Biol. Reprod. 95, 16, 1–10 (2016).

7. Evertts, A. G. et al. H4K20 methylation regulates quiescence and chromatin compaction. Mol. Biol. Cell 24, 3025–3037 (2013).

8. Swygert, S. G. et al. Condensin-Dependent Chromatin Compaction Represses Transcription Globally during Quiescence. Mol. Cell 73, 533–546.e4 (2019).

9. Coller, H. A., Sang, L. & Roberts, J. M. A New Description of Cellular Quiescence. PLOS Biol. 4, e83 (2006).

10. Bjornson, C. R. R. et al. Notch Signaling Is Necessary to Maintain Quiescence in Adult Muscle Stem Cells. Stem Cells 30, 232–242 (2012).

11. Sang, L., Coller, H. A. & Roberts, J. M. Control of the Reversibility of Cellular Quiescence by the Transcriptional Repressor HES1. Science 321, 1095–1100 (2008).

12. Valcourt, J. R. et al. Staying alive: Metabolic adaptations to quiescence. Cell Cycle 11, 1680–1696 (2012).

13. Legesse-Miller, A. et al. Quiescent fibroblasts are protected from proteasome inhibition–mediated toxicity. Mol. Biol. Cell 23, 3566–3581 (2012).

14. Tang, A. H. & Rando, T. A. Induction of autophagy supports the bioenergetic demands of quiescent muscle stem cell activation. EMBO J. 33, 2782–2797 (2014).

15. Lemons, J. M. S. et al. Quiescent Fibroblasts Exhibit High Metabolic Activity. PLOS Biol. 8, e1000514 (2010).

16. Leeman, D. S. et al. Lysosome activation clears aggregates and enhances quiescent neural stem cell activation during aging. Science 359, 1277–1283 (2018).

17. Pardee, A. B. A Restriction Point for Control of Normal Animal Cell Proliferation. Proc. Natl. Acad. Sci. 71, 1286–1290 (1974).

18. Polyak, K. et al. Cloning of p27Kip1, a cyclin-dependent kinase inhibitor and a potential mediator of extracellular antimitogenic signals. Cell 78, 59–66 (1994).

19. Criscione, S. W. et al. Reorganization of chromosome architecture in replicative cellular senescence. Sci. Adv. 2, e1500882 (2016).

20. Young, C. P. et al. Distinct histone methylation and transcription profiles are established during the development of cellular quiescence in yeast. BMC Genomics 18, 107 (2017).

21. Liu, L. et al. Chromatin Modifications as Determinants of Muscle Stem Cell Quiescence and Chronological Aging. Cell Rep. 4, 189–204 (2013).

22. Bonitto, K., Sarathy, K., Atai, K., Mitra, M. & Coller, H. A. Is There a Histone Code for Cellular Quiescence? Front. Cell Dev. Biol. 9, (2021).

23. Gray, J. V. et al. “Sleeping Beauty”: Quiescence in Saccharomyces cerevisiae. Microbiol. Mol. Biol. Rev. 68, 187–206 (2004).

24. Mitra, M. et al. Alternative polyadenylation factors link cell cycle to migration. Genome Biol. 19, 176 (2018).

25. Hu, K. et al. A Reversible Neural Stem Cell Quiescence and Activation Culture System for Metabolic Study. Cell Transplant. 33, 09636897241259723 (2024).

26. Venezia, T. A. et al. Molecular Signatures of Proliferation and Quiescence in Hematopoietic Stem Cells. PLOS Biol. 2, e301 (2004).

27. Ong, C.-T. & Corces, V. G. CTCF: an architectural protein bridging genome topology and function. Nat. Rev. Genet. 15, 234–246 (2014).

28. Narita, T. et al. Disentangling the architectural and non-architectural functions of CTCF and cohesin in gene regulation. Nat. Genet. 57, 3137–3151 (2025).

29. Kourmouli, N. et al. Heterochromatin and tri-methylated lysine 20 of histone H4 in animals. J. Cell Sci. 117, 2491–2501 (2004).

30. Schotta, G. et al. A silencing pathway to induce H3-K9 and H4-K20 trimethylation at constitutive heterochromatin. Genes Dev. 18, 1251–1262 (2004).

31. Stender, J. D. et al. Control of Proinflammatory Gene Programs by Regulated Trimethylation and Demethylation of Histone H4K20. Mol. Cell 48, 28–38 (2012).

32. Bierhoff, H. et al. Quiescence-Induced LncRNAs Trigger H4K20 Trimethylation and Transcriptional Silencing. Mol. Cell 54, 675–682 (2014).

33. Zhao, Z., Dammert, M. A., Grummt, I. & Bierhoff, H. lncRNA-Induced Nucleosome Repositioning Reinforces Transcriptional Repression of rRNA Genes upon Hypotonic Stress. Cell Rep. 14, 1876–1882 (2016).

34. Iyer, V. R. et al. The Transcriptional Program in the Response of Human Fibroblasts to Serum. Science 283, 83–87 (1999).

35. Kurup, J. T., Han, Z., Jin, W. & Kidder, B. L. H4K20me3 methyltransferase SUV420H2 shapes the chromatin landscape of pluripotent embryonic stem cells. Development 147, dev188516 (2020).

36. Layer, R. M. et al. GIGGLE: a search engine for large-scale integrated genome analysis. Nat. Methods 15, 123–126 (2018).

37. Singh, R. & Berger, B. Deciphering the species-level structure of topologically associating domains. 2021.10.28.466333 Preprint at 10.1101/2021.10.28.466333 (2021).

38. Krietenstein, N. et al. Ultrastructural Details of Mammalian Chromosome Architecture. Mol. Cell 78, 554–565.e7 (2020).

39. Bromberg, K. D. et al. The SUV4-20 inhibitor A-196 verifies a role for epigenetics in genomic integrity. Nat. Chem. Biol. 13, 317–324 (2017).

40. Abini-Agbomson, S. et al. Catalytic and non-catalytic mechanisms of histone H4 lysine 20 methyltransferase SUV420H1. Mol. Cell 83, 2872–2883.e7 (2023).

41. Secchia, S. et al. Starvation activates ECM-remodeling gene transcription and putative enhancers in fibroblasts despite inducing quiescence. Cell Rep. 44, (2025).

42. Hyle, J. et al. Acute depletion of CTCF directly affects MYC regulation through loss of enhancer–promoter looping. Nucleic Acids Res. 47, 6699–6713 (2019).

43. Schotta, G. et al. A chromatin-wide transition to H4K20 monomethylation impairs genome integrity and programmed DNA rearrangements in the mouse. Genes Dev. 22, 2048–2061 (2008).

44. Nelson, D. M. et al. Mapping H4K20me3 onto the chromatin landscape of senescent cells indicates a function in control of cell senescence and tumor suppression through preservation of genetic and epigenetic stability. Genome Biol. 17, 158 (2016).

45. Rao, S. S. P. et al. A 3D Map of the Human Genome at Kilobase Resolution Reveals Principles of Chromatin Looping. Cell 159, 1665–1680 (2014).

46. Tam, P. L. F., Cheung, M. F., Chan, L. Y. & Leung, D. Cell-type differential targeting of SETDB1 prevents aberrant CTCF binding, chromatin looping, and cis-regulatory interactions. Nat. Commun. 15, 15 (2024).

47. Strickfaden, H., Missiaen, K., Hendzel, M. J. & Underhill, D. A. KMT5C displays robust retention and liquid-like behavior in phase separated heterochromatin. 776625 Preprint at 10.1101/776625 (2019).

48. Ren, W. et al. DNMT1 reads heterochromatic H4K20me3 to reinforce LINE-1 DNA methylation. Nat. Commun. 12, 2490 (2021).

49. Hashimoto, H. et al. Structural Basis for the Versatile and Methylation-Dependent Binding of CTCF to DNA. Mol. Cell 66, 711–720.e3 (2017).

50. Serna-Pujol, N. et al. Coordinated changes in gene expression, H1 variant distribution and genome 3D conformation in response to H1 depletion. Nucleic Acids Res. 50, 3892–3910 (2022).

51. Guerra-Calderas, L. et al. CTCF-KDM4A complex correlates with histone modifications that negatively regulate CHD5 gene expression in cancer cell lines. Oncotarget 9, 17028–17042 (2018).

52. Hahn, M. et al. Suv4-20h2 mediates chromatin compaction and is important for cohesin recruitment to heterochromatin. Genes Dev. 27, 859–872 (2013).

53. Palikyras, S. et al. Senescent cells cluster CTCF on nuclear speckles to instruct an alternative splicing program. *Nat*. Aging 1–19 (2026) doi:10.1038/s43587-026-01171-6.

54. Criscione, S. W. et al. Reorganization of chromosome architecture in replicative cellular senescence. Sci. Adv. 2, e1500882 (2016).

55. Ravikanth, M., Soujanya, P., Manjunath, K., Saraswathi, T. & Ramachandran, C. Heterogenecity of fibroblasts. J. Oral Maxillofac. Pathol. JOMFP 15, 247–250 (2011).

56. Rawlings, J. S., Gatzka, M., Thomas, P. G. & Ihle, J. N. Chromatin condensation via the condensin II complex is required for peripheral T-cell quiescence. EMBO J. 30, 263–276 (2011).

57. Neumann, F. R. & Nurse, P. Nuclear size control in fission yeast. J. Cell Biol. 179, 593–600 (2007).

58. Jorgensen, P. et al. The Size of the Nucleus Increases as Yeast Cells Grow. Mol. Biol. Cell 18, 3523–3532 (2007).

59. Mitra, M., Batista, S. L. & Coller, H. A. Transcription factor networks in cellular quiescence. Nat. Cell Biol. 27, 14–27 (2025).

60. Wongtawan, T., Taylor, J. E., Lawson, K. A., Wilmut, I. & Pennings, S. Histone H4K20me3 and HP1α are late heterochromatin markers in development, but present in undifferentiated embryonic stem cells. J. Cell Sci. 124, 1878–1890 (2011).

61. Chomiak, A. A. et al. Nde1 is required for heterochromatin compaction and stability in neocortical neurons. iScience 25, 104354 (2022).

62. Gregor, A. et al. De Novo Mutations in the Genome Organizer CTCF Cause Intellectual Disability. Am. J. Hum. Genet. 93, 124–131 (2013).

63. Schindelin, J. et al. Fiji: an open-source platform for biological-image analysis. Nat. Methods 9, 676–682 (2012).

64. Frankish, A. et al. GENCODE reference annotation for the human and mouse genomes. Nucleic Acids Res. 47, D766–D773 (2019).

65. Salmon provides fast and bias-aware quantification of transcript expression | Nature Methods. https://www.nature.com/articles/nmeth.4197.

66. Soneson, C., Love, M. I. & Robinson, M. D. Differential analyses for RNA-seq: transcript-level estimates improve gene-level inferences. Preprint at 10.12688/f1000research.7563.2 (2016).

67. Love, M. I., Huber, W. & Anders, S. Moderated estimation of fold change and dispersion for RNA-seq data with DESeq2. Genome Biol. 15, 550 (2014).

68. Wu, T. et al. clusterProfiler 4.0: A universal enrichment tool for interpreting omics data. The Innovation 2, 100141 (2021).

69. Joung, J. et al. A transcription factor atlas of directed differentiation. Cell 186, 209–229.e26 (2023).

70. Kaya-Okur, H. S., Janssens, D. H., Henikoff, J. G., Ahmad, K. & Henikoff, S. Efficient low-cost chromatin profiling with CUT&Tag. Nat. Protoc. 15, 3264–3283 (2020).

71. Zheng, Y., Ahmad, K. & Henikoff, S. CUT&Tag Data Processing and Analysis Tutorial. https://www.protocols.io/view/cut-amp-tag-data-processing-and-analysis-tutorial-bjk2kkye (2020).

72. Langmead, B. & Salzberg, S. L. Fast gapped-read alignment with Bowtie 2. Nat. Methods 9, 357–359 (2012).

73. Picard Tools - By Broad Institute. http://broadinstitute.github.io/picard/.

74. Amemiya, H. M., Kundaje, A. & Boyle, A. P. The ENCODE Blacklist: Identification of Problematic Regions of the Genome. Sci. Rep. 9, 9354 (2019).

75. Meers, M. P., Tenenbaum, D. & Henikoff, S. Peak calling by Sparse Enrichment Analysis for CUT&RUN chromatin profiling. Epigenetics Chromatin 12, 42 (2019).

76. Stark, R. & Brown, G. DiffBind: Differential binding analysis of ChIP-Seq peak data.

77. Zheng, R. et al. Cistrome Data Browser: expanded datasets and new tools for gene regulatory analysis. Nucleic Acids Res. 47, D729–D735 (2019).

78. Bailey, T. L. & Grant, C. E. SEA: Simple Enrichment Analysis of motifs. 2021.08.23.457422 Preprint at 10.1101/2021.08.23.457422 (2021).

79. Zheng, B., Zhang, Z., Black, C. M., Crombrugghe, B. de & Denton, C. P. Ligand-Dependent Genetic Recombination in Fibroblasts: A Potentially Powerful Technique for Investigating Gene Function in Fibrosis. Am. J. Pathol. 160, 1609–1617 (2002).

80. Jozefczuk, J., Drews, K. & Adjaye, J. Preparation of Mouse Embryonic Fibroblast Cells Suitable for Culturing Human Embryonic and Induced Pluripotent Stem Cells. J. Vis. Exp. JoVE e3854 (2012) doi:10.3791/3854.

